# To mix or to de-mix? : Cation-π suppresses heterogeneity in local structure and dynamics of heterotypic Condensates

**DOI:** 10.64898/2026.10.10.758149

**Authors:** Sudhriti Roy, Deepak Sharma, Milan Kumar Hazra

## Abstract

Heterotypic biomolecular condensates can exhibit pronounced spatial and dynamic heterogeneity because chemically distinct components compete between self- and cross-association. Here, we use coarse-grained simulations to investigate how cation–π interactions modulate this competition in condensates formed by peptides with systematic variation of hydrophobicity-to-charge ratios in binary peptide pairs. By independently tuning sequence heterogeneity, peptide composition, and cation–π interaction strength, we establish how an additional short-range interaction reshapes condensate stability, internal organization, and molecular dynamics. In the absence of cation–π interactions, highly heterogeneous peptide pairs form distinct core–shell architectures, with charge- and hydrophobicity-rich sequences preferentially occupying different regions of the condensate. Introducing cation–π interactions progressively enhances phase stability and promotes molecular mixing, with stronger interactions substantially increasing the critical temperature and condensate density. Energetic analysis reveals that cation–π interactions partially compensate for the loss of electrostatic and hydrophobic stabilization, providing an additional and tuneable source of cohesive interactions. This enhanced cross-association also suppresses spatial and dynamic heterogeneity, yielding more homogeneous peptide distributions, slower translational diffusion, and more uniform chain reconfiguration within the dense phase. Collectively, our results demonstrate that cation–π interactions provide a molecular mechanism for coupling condensate stability to spatial organization and internal dynamics, enabling sequence heterogeneity and composition to be translated into distinct condensate states.

## Introduction

Biomolecular condensates organize cellular components without membrane boundaries through the collective association of proteins, nucleic acids, and other biomolecules^1–4^. By concentrating selected molecules while permitting continuous exchange with the surrounding phase, condensates regulate diverse processes including RNA metabolism, transcription, signalling, stress responses, and intracellular organization^5^. Their dynamic nature arises from reversible and multivalent interactions whose collective behaviour is sensitive to molecular concentration, sequence, post-translational modification, and environmental conditions^1,4,5^. Condensates are therefore best viewed as dynamic molecular networks whose mesoscale organization and material properties emerge from interactions operating across multiple length and time scales^4,6^.

Intrinsically disordered proteins (IDPs) and intrinsically disordered regions (IDRs) are particularly effective condensate-forming molecules because their conformational flexibility enables numerous transient intermolecular contacts^7–10^. Unlike folded proteins, IDPs sample heterogeneous conformational ensembles and can dynamically expose interaction motifs involving charged, aromatic, hydrophobic, polar, and hydrogen-bonding residues^8,9,11–15^. Consequently, condensate behaviour depends not only on overall composition but also on the identity, valence, spacing, and patterning of interaction-prone residues.[9–12] This sequence dependence has motivated the concept of a molecular “grammar” of condensation, in which relatively subtle changes in primary sequence can alter phase behaviour, molecular partitioning, and material properties^4,11,16^.

The stickers-and-spacers framework provides a useful physical description of this sequence dependence^17,18^. Multivalent sticker residues or motifs form transient attractive contacts, while spacers regulate chain conformations, interaction accessibility, and network connectivity. Importantly, stickers can engage through multiple interaction classes, including electrostatic, hydrophobic, aromatic, cation–π, and hydrogen-bonding interactions^15,19–23^. Condensate properties therefore reflect the competition and cooperation among these interactions rather than a single dominant energetic contribution^24–28^. Indeed, computational and experimental studies have shown that both aromatic residue patterning and charge organization can strongly modulate phase behaviour and dense-phase material properties^15,29–34^.

These effects become particularly complex in heterotypic condensates, where chemically distinct molecular populations undergo simultaneous self- and cross-association. When self-interactions dominate, the components may segregate into distinct domains or core–shell-like organizations; favourable heterotypic interactions, in contrast, can promote molecular mixing^35,36^. The resulting architecture is therefore governed by the relative free energies of homotypic and heterotypic contacts. Molecular composition provides a further control parameter by changing the availability of self- and cross-interactions and can determine whether a component forms a continuous scaffold, preferentially occupies the condensate interior or interface, or remains distributed throughout the dense phase^4,35,36^.

Sequence heterogeneity offers a direct means of tuning this interaction balance. Two IDPs can have similar overall physicochemical properties while differing substantially in the distribution of hydrophobic, charged, or aromatic residues. Hydrophobic and aromatic residues can provide short-range cohesive interactions, whereas charged residues generate electrostatic attraction or repulsion depending on their partners^22,29,37^. Importantly, the distribution of interaction types between molecular species may be more consequential than their average abundance. A peptide enriched in charged residues may preferentially stabilize itself through electrostatic interactions, whereas a second peptide enriched in hydrophobic/aromatic residues may favour short-range contacts. Such interaction mismatch can generate spatial differentiation and component-specific dynamics within the condensate^30,38^.

Our recent work demonstrated that sequence-level heterogeneity can directly regulate stability and dynamics of heterotypic peptide condensates. Differential charge clustering between two polyampholyte populations enhanced condensate stability relative to homogeneous reference systems and generated pronounced differences in molecular diffusivity^30^. Increasing charge-patterning disparity amplified structural and dynamical heterogeneity, while the mixing ratio provided an additional means of tuning condensate stability and mobility. These findings established sequence heterogeneity and composition as independent control parameters for heterotypic condensate organization. However, this framework was dominated by long-range electrostatic interactions. Because biological condensates simultaneously contain multiple interaction types, an important unresolved question is how a short-range heterotypic interaction modifies condensate organization arising from sequence mismatch.

Cation–π interactions provide a particularly compelling mechanism for such coupling. They arise from favourable association between a positively charged group and the electron-rich face of an aromatic ring, combining electrostatic and aromatic contributions within a spatially localized contact^8,39^. Arginine and lysine on one side and phenylalanine, tyrosine, and tryptophan on the other are abundant in IDRs, making cation–π interactions natural candidates for coupling otherwise chemically distinct molecular populations^8,23,40^. Unlike long-range electrostatic interactions, cation–π contacts are short-ranged and sensitive to local geometry and residue accessibility, allowing them to function as localized cross-links within heterogeneous interaction networks^41^.

Experimental and computational studies support a role for cation–π interactions in disordered-protein assembly. In FUS, for example, interactions between arginine-rich and aromatic regions contribute to intermolecular association, while modification of arginine residues alters phase behaviour and material properties^40^. Studies of DDX4 and related systems further demonstrate that electrostatic, aromatic, and hydrophobic interactions collectively determine sequence-dependent phase behavior^29^. Importantly, these studies also show that cation–π interactions cannot be treated as an independent or universally dominant driving force. A recent study of arginine-rich and tyrosine-rich polypeptides found that cation–π interactions contribute to phase separation but that Coulombic interactions and tyrosine-mediated hydrogen bonding can make substantial contributions depending on molecular context.[29] Thus, the effect of cation–π interactions must be understood within the broader interaction network rather than as a simple additional attractive energy.

This consideration is particularly important for heterotypic condensates composed of chemically mismatched peptides. When one population is enriched in charged residues and the other in aromatic/hydrophobic residues, their preferred interaction environments differ: electrostatic association can dominate one population, whereas short-range hydrophobic/aromatic interactions stabilize the other. Such mismatch can favour spatial differentiation. Introducing cation–π interactions creates a direct cross-association pathway between these populations and may therefore reduce the thermodynamic advantage of segregation and promote molecular mixing. At the same time, stronger cross-association increases network connectivity and can constrain molecular motion. Thus, enhanced spatial mixing does not necessarily imply enhanced mobility; the same interaction can simultaneously homogenize composition and dynamically confine the constituent molecules.

Moreover, spatial and dynamical heterogeneity need not change in parallel. Two molecular populations may become spatially mixed while retaining different local mobilities if their interaction environments remain distinct, whereas sufficiently strong cross-association can generate a common interaction network and reduce both compositional and dynamical differences.[20,21,30] A complete description of heterotypic condensates therefore requires simultaneous consideration of thermodynamic stability, internal organization, molecular diffusion, intermolecular exchange, and conformational dynamics.

Here, we investigate how tuneable cation–π interactions reshape sequence-mismatched heterotypic peptide condensates using a systematic coarse-grained model. We consider two 40-residue disordered peptide populations with systematically varied hydrophobicity-to-charge ratios, generating three levels of sequence mismatch while maintaining comparable molecular size and average composition. The peptides are mixed at controlled ratios, and phenylalanine-mediated short-range hydrophobic interactions are combined with screened electrostatics. Cation–π interactions between positively charged and aromatic residues are then introduced with controlled strengths, allowing their contribution to be isolated without changing the underlying peptide sequences.

We examine the consequences of cation–π coupling at three interconnected levels: (i) condensate stability and density, to quantify its contribution to dense-phase stabilization; (ii) spatial organization and morphology, to determine whether cross-association promotes molecular mixing or preserves sequence-dependent segregation; and (iii) translational diffusion, neighbour exchange, and conformational reconfiguration, to establish how structural reorganization translates into molecular dynamics. Particular attention is given to component-specific dynamics, allowing global condensate stabilization to be distinguished from selective kinetic confinement.

We find that cation–π interactions act as more than an additional source of cohesive energy. In strongly sequence-mismatched systems, increasing cation–π coupling progressively overcomes the spatial differentiation generated by competing electrostatic and hydrophobic interactions and promotes a more homogeneous distribution of the two peptide populations. Concurrently, stronger cation–π coupling enhances condensate stability and density while reducing molecular diffusion and conformational reconfiguration. Thus, stronger heterotypic coupling can simultaneously promote spatial homogenization and dynamic confinement. Energetic analysis further reveals that cation–π interactions redistribute the relative contributions of electrostatic and hydrophobic contacts, demonstrating that their effect is cooperative rather than simply additive.

Together, these results establish a framework in which sequence mismatch drives molecular differentiation, composition regulates the availability of self- and cross-interactions, and cation–π interactions provide a tuneable pathway for coupling chemically distinct peptide populations. More broadly, our findings show that condensate stability alone does not determine internal organization or material behaviour: the same interaction that strengthens condensation can simultaneously reshape molecular partitioning, morphology, exchange kinetics, and conformational dynamics. Understanding this coupling between interaction chemistry and mesoscale material properties provides a route toward predictive control of multicomponent biomolecular condensates.

## Methods and Simulation Details

To elucidate phase behaviour of heterotypic condensates composed of two distinct sequence types we employ a C-alpha based coarse-grained model of peptides. Each peptide consists of 40 residues, with each residue represented by a single coarse-grained bead. To systematically investigate the effect of sequence heterogeneity on condensate organization, three pairs of sequences with different values of were considered. The heterogeneity has been driven by hydrophobicity to charge ratio as shown in Figure 1. Three Δϕ variants has been designed with peptide pairs having hydrophobicity to charged residue ratio ϕ=0.25 and ϕ=2.33 (Δϕ=2.08), ϕ=0.42 and ϕ=1.5 (Δϕ=1.08), ϕ=0.66 and ϕ=1.00 (Δϕ=0.34)

**Figure 1.**
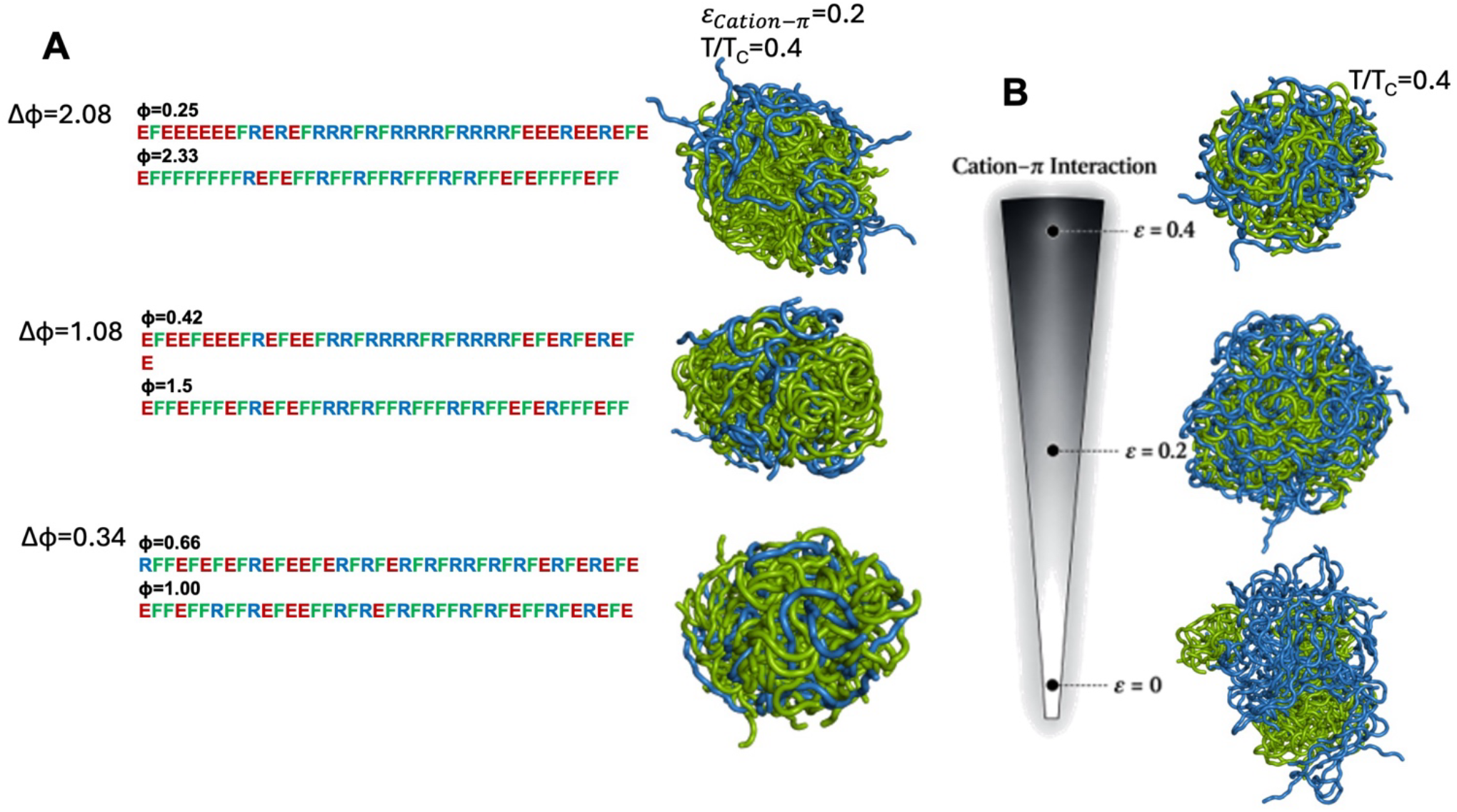
(A) Designed sequence pairs with differential hydrophobicity to charge ratios (Δϕ). Heterotypic condensates obtained in weak cation-π limit (ε_*Cation*−π_= 0. 2) at temperature T/T_C_=0.4 and at asymmetric mixing fraction *x*_*High*−*ϕ*_ *=* 0. 75 for Δϕ variants namely 2.08, 1.08 and 0.34. (B) Effect of cation-π interaction strength on mixing and dissolution of multiphases in condensate for peptide pairs for Δϕ=2.08 at *x*_*High*−*ϕ*_ *=* 0. 50. Blue and green cartoon represents low and high hydrophobicity/charge ratio in neutral peptides.

The coarse-grained model consists of bonded interactions that maintain the connectivity of the polymer chains and nonbonded interactions describing electrostatic, attractive, and excluded-volume interactions. The electrostatic interactions between charged residues are described using a screened Coulomb or Debye–Hückel potential^42^,

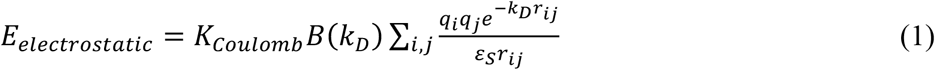

where *q*_*i*_ and *q*_*j*_ denote the charge of the *i*^th^ and j^th^ bead, *r*_*ij*_ denotes the inter-bead distance, ε_*s*_ is solvent dielectric constant, and *K*_*coulomb*_= *4π*ε_0_=332 kcal/mol. *B*(*k*_*D*_) is a function of solvent salt concentration and the radius (a) of ions produced by the dissociation of the salt, and can be expressed as 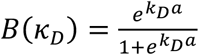. The Debye-Hückel electrostatic interactions of an ion pair act over a length scale of the order of *k*^*−*1^, which is called the Debye screening length. *k*_*D*_ is related to ionic strength as,

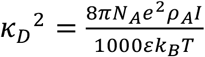

where *N*_*A*_ is the Avogadro number, *e* is the charge of an electron, ρ_*A*_ is the solvent density, *I* denotes solvent ionic strength, *k*_*B*_ is the Boltzmann constant, and T is the temperature. To avoid overlap among beads, a steep repulsion interaction has been defined as 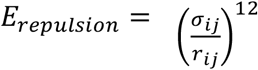 with σ_*ij*_=4 Å. Hydrophobic PHE residues interact through a short-range 12-10 dispersion interaction

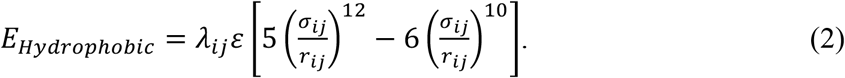

Phe–Phe interactions were described by a short-range potential with ε = 0.2 kcal mol^−1^, that reproduces experimentally observed dimensions of intrinsically disordered peptides^32^. Langevin dynamics simulations were performed for 100 binary polymers over a range of mixing fractions and subcritical temperatures at an ionic strength of 0.04 M using an implicit solvent. Two independent trajectories of 10^7^ integration steps were generated for each condition, and equilibrated largest condensates identified through cluster analysis were used for all structural, thermodynamic, and dynamical analyses.

To investigate the role of cation-π interactions in controlling the organization of heterotypic condensates, we extend the baseline model by introducing an additional attractive interaction between positively charged and aromatic residues. The cation-π interaction is represented by a the same 12-10 potential as described earlier, the strength of cation-π interactions we varied having a weak interaction limit (ε_*Cation–π*_ = 0.2) and a strong interaction limit (ε_*Cation–π*_ = 0.4) and has been compared with the simulations switching off the cation-π interactions (ε_*Cation–π*_ = 0.0).

Langevin dynamics simulations were performed for each combination of sequence pair, mixing fraction, and interaction strength. The polymers were initially placed in a cubic simulation box extending from −150 to 150 Å along each Cartesian direction, corresponding to a box size of 300×300 × 300 Å^3^. An implicit salt concentration of 0.04 M was used for all simulations. Each system was propagated for a total of 8×10^6^ stochastic dynamics steps. Configurations were recorded every 500 integration steps, resulting in 16,000 configurations for each simulation. The first 8,000 configurations were discarded as the equilibration period, while the remaining 8,000 configurations were used for subsequent structural and thermodynamic analyses. Two independent copies of simulations has been performed for each system.

The equilibrated trajectories were analysed using cluster-based methods to identify the condensed phase and the largest droplet. Neighbour list of each peptide has been defined with a centre of mass (COM) distance cut-off 30 Å and having at least 5 inter chain contact beads. Breadth-first search algorithm has been used to find the largest cluster by identifying connected COMs of peptides in a graph. The structural and energetic properties of the condensates were evaluated as a function of sequence heterogeneity and cation-π interaction strength. In particular, comparison of systems with a spectrum of cation-π strength at different mixing fractions allows us to determine how the presence and strength of cation-π interactions influence the spatial organization and mixing/de-mixing behaviour of heterotypic condensates. This framework enables us to systematically disentangle the effects of sequence heterogeneity, composition, and cation-π cross interactions on the phase behaviour of the condensates.

## Results and Discussion

### Phase Diagram and Stability of Condensates

To evaluate how cation-π interactions affect condensate stability, we have computed phase diagrams for designed systems across weak (*E*_*Cation −*π_ *=* 0.2 kcal/mol) and strong (*E*_*Cation −*π_ *=* 0.4 kcal/mol)^41^ interaction regimes while varying peptide mixing fractions (*x*_*High −ϕ*_) and differential hydrophobic to charged residue ratio differences among pair peptides (Δϕ).Figure 1 (panels A to C showcases phase diagrams for Δϕ variant 2.08 at mixing fraction respectively *x*_*High −ϕ*_ =0.25, 0.50 and 0.75. Convergence of density in condensate phase has been shown from independent simulation runs at T/T_C_ =0.7 for *x*_*High −ϕ*_ =0.50 for all Δϕ variants at different cation-π strengths (Figure S1).A significant enhancement in droplet phase stability has been observed upon transitioning from a weak to a strong cation–π interaction regime.

At maximum sequence heterogeneity (Δϕ=2.08) and charge-dominated composition (*x*_*High−ϕ*_ = 0.25), condensates lacking cation-π interactions exhibit a critical reduced temperature of T_C_=0.8. Introducing weaker cation-π interactions elevates T_C_ by 25% (T_C_=1.0) and increases the condensate density by 1.3-fold near criticality (T/T_C_=0.8). At the strong cation-π limit, T_C_ increases by 50% (T_C_=1.2), driving a nearly 3-fold enhancement in condensate density (Figures 2A and 4A,B). For balanced sequence mixing fraction (*x*_*High−ϕ*_ *=* 0.50), weak cation-π interactions increase T_C_ by 30% and condensate density by 1.5-fold, whereas strong cation-π interactions yield a 60% enhancement in T_C_ and a 1.56-fold density increase at T/T_C_=0.8 (Figures 2B and 4A,B). In condensates enriched with sequences having high-ϕ (*x*_*High−ϕ*_ = 0.75), weak and strong cation-π coupling enhance T_C_ by 40% and 75%, respectively. While density variation remains minor under weak coupling, strong cation--π interactions increase condensate density by 1.56 times (Figures 2C and 4 A,B).

**Figure 2.**
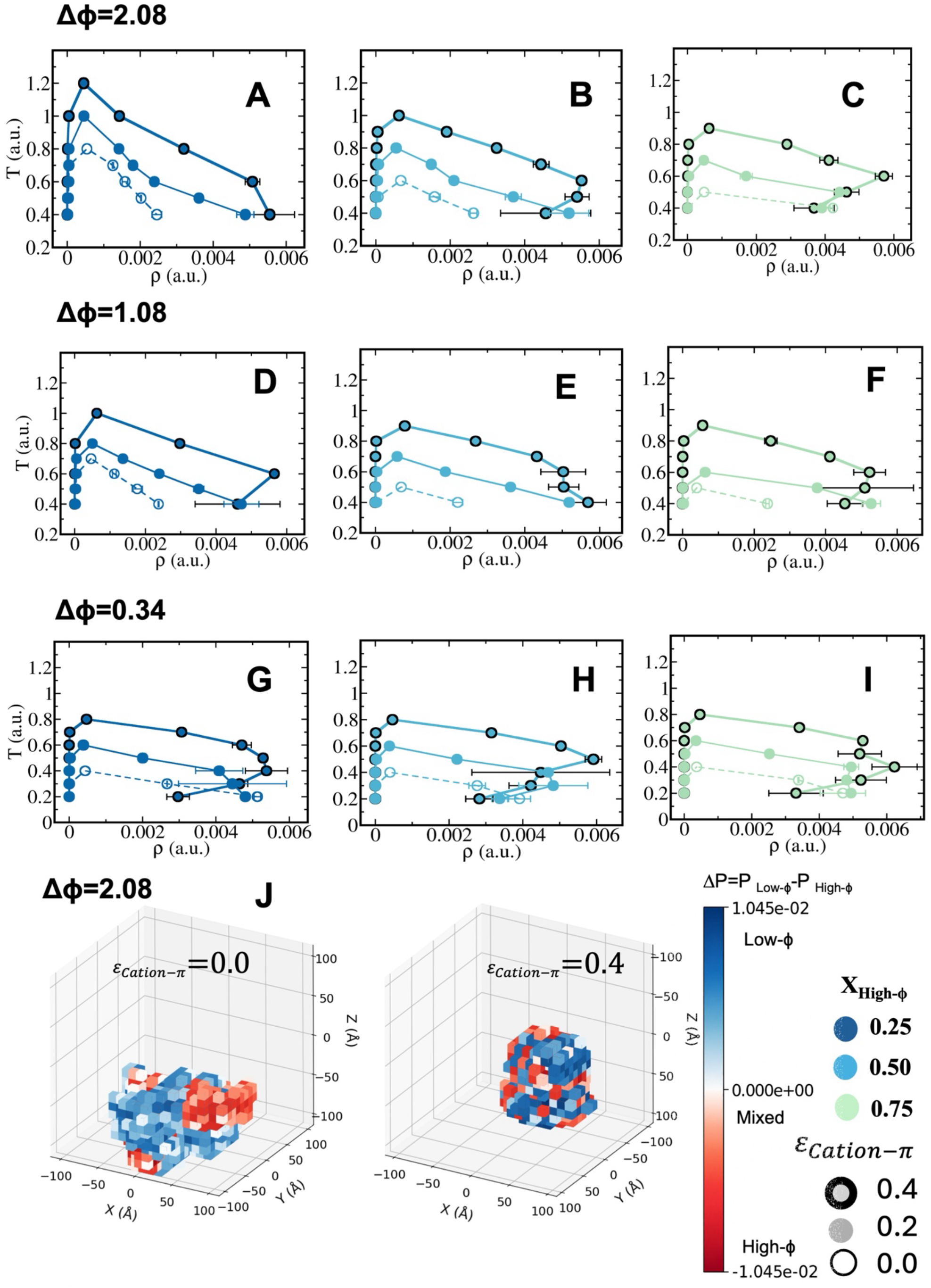
Coexistence curves denoting stability of condensates obtained peptide pairs with differential hydrophobicity to charge ratios (A-C) Δϕ=2.08, (D-F) Δϕ=1.08 and (G-I) Δϕ=0.34. Panels (A, D and G) respectively showcase phase diagrams at mixing fraction of peptides *x*_*High*−*ϕ*_ *=* 0. 25; panels B,E and H depict phase diagrams at *x*_*High*−*ϕ*_ *=* 0. 50 and panels C,F and I showcase phase diagrams at *x*_*High*−*ϕ*_ *=* 0. 75. In each panel, phase diagrams has been compared along cation-π strength ε_*Cation*−π_= 0. 0 (void symbol), ε_*Cation*−π_= 0. 2 *kcal*/*mol* (filled symbol) and ε_*Cation*−π_= 0. 4 *kcal*/*mol* (filled symbol with outer lines). Colour codes represent mixing fractions (*X*_*High*−*ϕ*_). (J) Spatial probability difference ΔP= *P*_*Low*−*ϕ*_ − *P*_*High*−*ϕ*_ in local grids on condensate at equimolar composition of peptides at highest sequence heterogeneity Δϕ=2.08 at ε_*Cation*−π_= 0. 0 *and* 0. 4. Red represents grids are filled with sequences with high-ϕ peptides and blue represents grids are filled with sequences with low-ϕ peptides. While extreme sequence heterogeneity de-mixes condensates into multiphases at ε_*Cation*−π_= 0. 0, the same extremity causes mixing at enhanced cation-π strengths.

At intermediate sequence heterogeneity (Δϕ= 1.08), the enhancement of the critical temperature is noticeably lower at asymmetric mixing fractions (*x*_*High−ϕ*_ = 0.25 a*nd* 0.75) compared to the highly heterogeneous regime of sequences (Δϕ= 2.08). This reduced thermal stabilization originates from fewer available cross-cation-π interactions between the asymmetric sequence pairs. Specifically, weak cation-π coupling yields only a ~20% increase in T_C_ relative to the cation-π-free baseline. Under strong cation-π coupling, T_C_ elevates by 40% at *x*_*High−ϕ*_ = 0.25 and 75% at *x*_*High−ϕ*_ = 0.75 (Figures 2D-F and 4A,B).

In contrast, the equimolar composition (*x*_*High−ϕ*_ = 0.50) exhibits a stronger criticality enhancement—increasing T_C_ by 40% at weak coupling and 80% at strong coupling—owing to equal peptide contributions and optimal sequence complementarity. Near criticality (T/T_C_=0.8), condensate density increases by 2-fold at *x*_*High−ϕ*_ = 0.25 across both weak and strong limits, whereas systems having abundance of sequences with high-ϕ achieve a 1.75-to 2.0-fold density amplification.

At vanishing heterogeneity of two sequences, we have observed almost no variation of T_C_ enhancement along mixing fraction having an increase of 50% at weak cation-π limit and an absolute 100% enhancement at strong cation-π limit. Density of the condensate is nearly 1.5-2 times higher than condensates without cation-π interactions along mixing fractions (Figures 2G-I and 4A,B).

The modulation of internal condensate heterogeneity by cation-π interactions is depicted in Figure 2J. To characterize the spatial organization within the dense phase, a grid-based framework was used to evaluate spatial probability density of peptides residing in the largest cluster in every 3D grid. Specifically, three-dimensional heatmaps representing the differential probability distributions at each grid of low and high-peptides

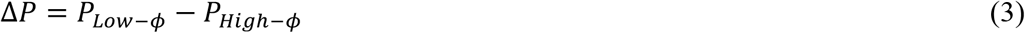

at equimolar fraction (*x*_*High−ϕ*_ = 0.50) highlighting the spatial heterogeneity within condensate phase. While cation-π interactions are absent, largest cluster showcase significant core-shell like architecture with high-ϕ peptides occupy surface of the largest cluster and peptides with dominant charged residues form the core of the droplet, at dominant cation-π interactions peptides mix in condensate uniformly and create a homogeneous mixture (Figure 2J and Figure S2). In addition we have showcased the effect of mixing fraction on condensate heterogeneity at different cation-π strengths for Δϕ=2.08 having highest sequence pair heterogeneity (Figure S2). It is observable that significant core-shell like architecture with preserved regions of low and high hydrophobic peptides has been observed in largest clusters in absence of cation-π interactions. While at low *x*_*High−ϕ*_, hydrophobic peptides occupy the surface and condensate core is being formed by charged residue rich peptides, at higher *x*_*High−ϕ*_, the opposite orientation prevails (Figure. S2).

### Energetic basis of Stability

Total energetic gain of the condensate along temperature (scaled to criticality) has been shown in Figure S3. Total energy that each peptide faces in a condensate varies with sequence pair hydrophobicity variation which even alters the charge density of the sequences (Figure S3). Total energetic gain per peptide in largest cluster is invariant to mixing fraction at a particular Δϕ. To dissect the energetic gain through different interactions we have segregated electrostatic, hydrophobic and cation-π components of the energetic stabilization along mixing fraction at T/T_C_=0.4 (Figure 3, S4). Interactions among low-ϕ peptides (P1-P1), high-ϕ peptides (P2-P2) and cross interactions among low-and high-ϕ peptides has been shown at T/TC=0.4 for all Δϕ and mixing fractions (Figure S5). Along enrichment of sequences with enhanced hydrophobicity to charge ratio, electrostatic interactions diminish and hydrophobic ones emerge linearly. It is interesting to note the depletion of hydrophobic interaction mediated stabilization at *x*_*High−ϕ*_ *=* 0.50 − 0.75 for Δϕ=2.08 (strongest dissimilarity of the sequence pairs leading to highest commitment to cation-π interactions) once we switch cation-π interactions. This indicates a balancing act of diminishing F-F interactions in terms of gaining R-F cross cation-π ones. At *x*_*High−ϕ*_ *=* 0.25, we observed 15 kcal/mol electrostatic stabilization of the condensate without any cation-π interactions decreases monotonically to 7 kcal/mol at highest enrichment of sequences with dominating hydrophobicity (Figure 3A). Hydrophobic interactions showcase a drop from 14kcal/mol to 12 kcal/mol at highest enrichment of sequences with hydrophobic nature once we switch on cation-π interactions (Figure 3B). A linear 3-fold diminishment is evident in hydrophobic interactions in largest cluster along enhancement of sequences with enhanced charged residues over hydrophobic ones.

**Figure 3.**
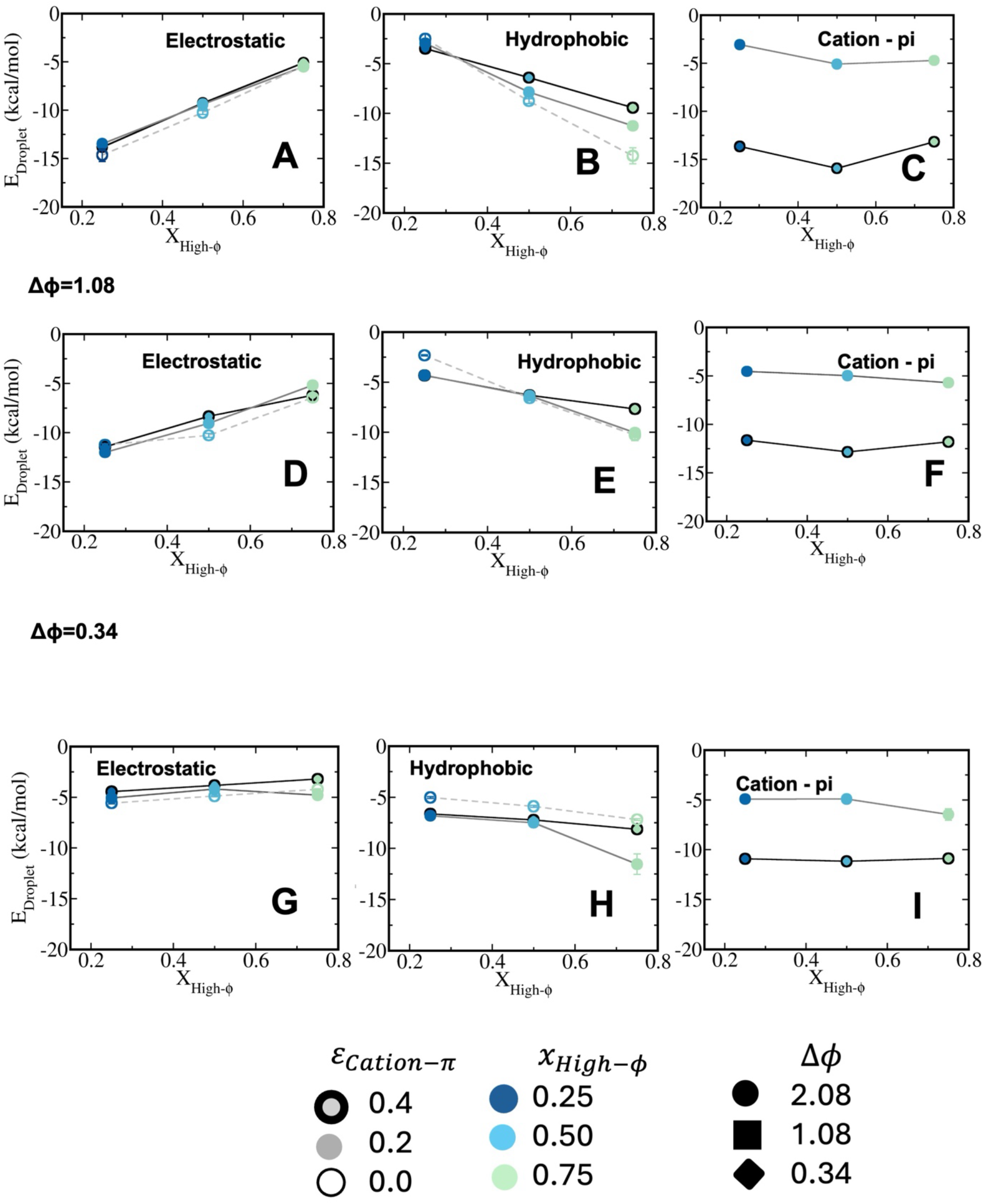
Long-range electrostatic, short-range hydrophobic and cation-π energy contributions in stabilizing heterotypic condensates for differential sequence hydrophobicity to charge ratio (A-C) Δϕ = 2.08, (D-F) Δϕ = 1.08 and (G-I) Δϕ = 0.34. Colour codes represent mixing fractions (*X*_*High*−ϕ_). Panels A,B and C; D,E and F and G,H and I represent long-range electrostatics, short range hydrophobic and cation-π energy contributions per peptide to total energy (experienced by each peptide on average) of the droplet phase along mixing fractions for Δϕ = 2.08; 1.08 and 0.34 variants respectively. In each panel, energy components have been compared along cation-π strength ε_*Cation*−π_= 0. 0 (void symbol), ε_*Cation*−π_= 0. 2 *kcal*/*mol* (filled symbol) and ε_*Cation*−π_= 0. 4 *kcal*/*mol* (filled symbol with outer lines).

Meanwhile cation-π interactions showcase almost no variations along *x*_*High−ϕ*_ but showcasing a shallow minimum at *x*_*High−ϕ*_ *=* 0.50 evidently having enhanced cation-π at equal mixing of the dissimilar sequences (Figure 3C). At lower cation-π limit the stabilization pertained from cation-π interactions ranges nearly 3-5 kcal/mol whereas at strong cation-π limit the same enhances up to 13-16 kcal/mol (nearly 3-4 folds).

At intermediate dissimilarity of sequence-pairs, we have observed 2kcal/mol drop in electrostatic interactions due to cation-π switching at *x*_*High−ϕ*_ *=* 0.50. Along enrichment of sequences with dominating hydrophobic interactions (*x*_*High−ϕ*_), electrostatics drops 2-fold and vice-versa for hydrophobic interactions (Figure 3D-E). At weak interacting limit, cation-π contributes 5 kcal/mol and at strong interaction limit the same ranges up to 12-14 kcal/mol irrespective of mixing (Figure 3F).

In absence of sequence heterogeneity (Figure 3G-I), interactions are nearly invariant to mixing fraction and even independent of varying cation-π strength. Electrostatic energy contributes nearly 5-8 kcal/mol, hydrophobic interactions contribute 5-11 kcal/mol independent upon mixing fraction. Hydrophobic interactions showcase a synergistic effect due to cooperativity at extreme *x*_*High−ϕ*_. Cation-π interaction is nearly 5 kcal/mol at weaker cation-π limit and nearly 11 kcal/mol at strong cation-π systems.

To evaluate the energetic stability underneath phase diagrams along sequence heterogeneity and cation-π strength, we have evaluated difference in energy each peptide faces in presence and in absence of cation-π interactions as sequence heterogeneity grows. At weaker cation-π limit the stabilization is independent of sequence heterogeneity and nearly 3-4kcal/mol in different mixing fractions of peptides. At stronger cation-π limit we observer a linear dependence of stability on sequence heterogeneity independent of mixing fractions ranging nearly 9 kcal/mol at weaker sequence heterogeneity (Δϕ=0.34) to 14 kcal/mol at strong sequence heterogeneity (Δϕ=2.08) (Figure 4C).

**Figure 4.**
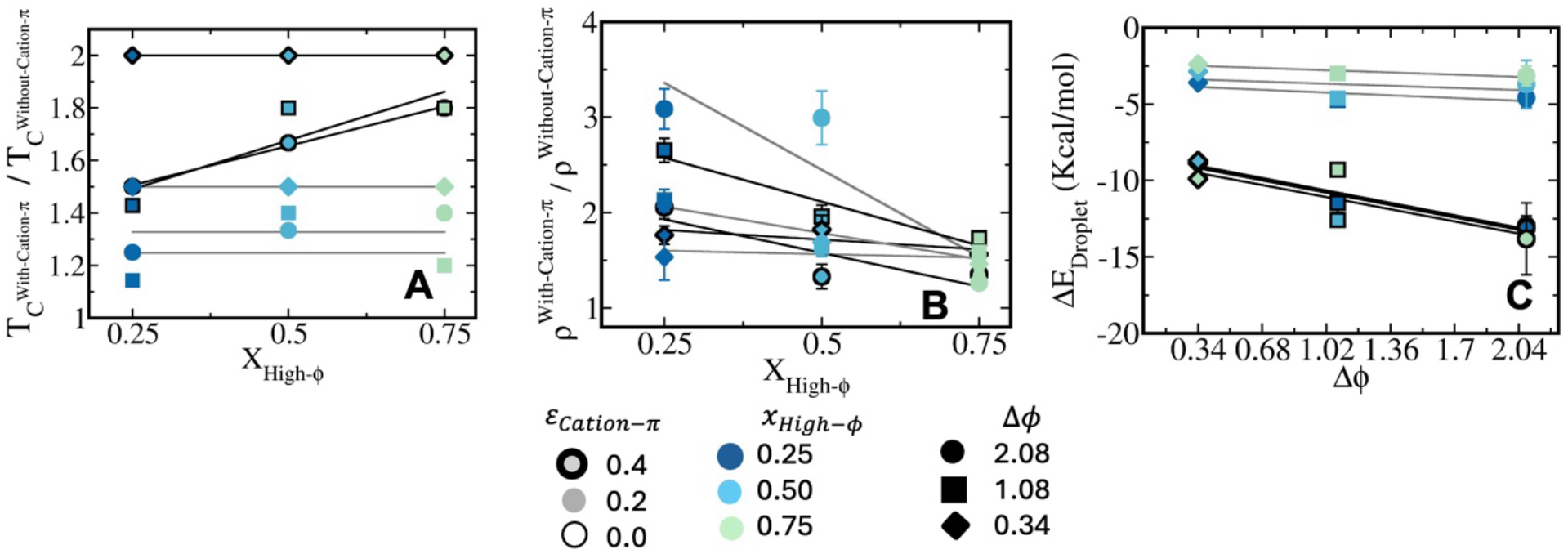
Comparative analysis of phase stability of heterotypic condensates with and without cation-π contribution along mixing fraction, sequence heterogeneity (A) Variation of critical temperature of the condensates with cation-π contribution with respect to condensates without any cation-π effect along mixing fraction at weaker (ε_*Cation*−π_= 0. 2 *kcal*/*mol*, filled symbols fitted with grey lines) and strong cation-π limit (ε_*Cation*−π_= 0. 4 *kcal*/*mol*, filled symbols with outer lines and fitted to black). (B) Variation of density of the condensates with cation-π contribution with respect to condensates without any cation-π effect along mixing fraction at weaker (ε_*Cation*−π_= 0. 2 *kcal*/*mol*, filled symbols fitted with grey lines) and strong cation-π limit (ε_*Cation*−π_= 0. 4 *kcal*/*mol*, filled symbol with outer lines and fitted to black). (C) Energetic stabilization of dense phase at weak and strong cation-π limit along sequence heterogeneity. Colour codes represent mixing fractions (*X*_*High*−ϕ_). Circles, squares and diamond symbols denote different Δϕ variants respectively. Gray lines show fits for cation-π strength ε_*Cation*−π_= 0. 2 *kcal*/*mol* and black lines show fits for ε_*Cation*−π_= 0. 4 *kcal*/*mol*.

### Translational Diffusivity and Reconfiguration Dynamics in Condensate Phase

In order to elucidate the effect of cation-π interactions on internal dynamics of condensate phase, we have computed mean squared displacements of peptides with different hydrophobicity-to-charge ratio in condensates. Mean squared displacements (MSD) has been calculated from trajectories as

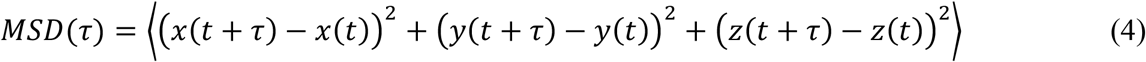

Convergence of mean squared displacements have been shown from independent trajectories in Figure S6 for representative systems at T/T_C_=0.4 and *X*_*High−ϕ*_= 0.50. Diffusion coefficients have been obtained from MSDs as slopes from a linear fit of the diffusive regime following the relation *MSD*(τ)= *6D*τ. In addition, we have quantified chain reconfigurational dynamics in condensate and bulk through time-correlation function of end-to-end distance of chains

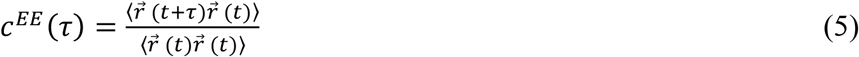

where 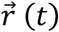 represents end-to-end distance vector of peptides. The correlation function has been fitted with a best fit bi-exponential function to evaluate the average timescale of reconfiguration. Convergence of time correlation functions has been shown from independent runs at T/T_C_=0.4 and *X*_*High−ϕ*_ =0.50(Figure S7).

Figure 5(A-C) depicts the ratio of 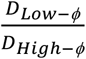 along mixing fraction for different Δϕ variants (panels A-C). 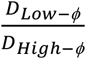 showcases a prominent enhancement along mixing fraction of sequences with enriched hydrophobic residues and low charge density. Diffusivity of low-ϕ peptide increases up to 2.5 times with respect to high-ϕ ones at increased *x*_*High−ϕ*_ for Δϕ=2.08 and 1.08 variant (Figure 5 A-B). Translational diffusivity is correlated to chain reconfigurational dynamics in condensates. At extreme sequence heterogeneity, average reconfiguration timescale of high-ϕ peptides is nearly 0.5-2 times of low-ϕ peptides and significantly dependent upon mixing fraction in absence of cation-π. At a lower mixing fraction (*x*_*High−ϕ*_), peptides with high-ϕ demonstrate a quicker reconfigurational time than low-ϕ ones, indicating that a dense core of droplet is formed by low-ϕ peptides and a shell by high-ϕ ones (Figure 2J and Figure S2). (Figure 2J and Figure S2). High-ϕ ones form droplet core and showcase significantly slow reconfigurational dynamics at higher *x*_*High−ϕ*_ and low-ϕ ones occupy surface of the droplet (Figure S2). In presence of cation-π interactions high-ϕ peptides showcase uniformly 1.5 times slower reconfigurational timescale than low-ϕ ones and is independent of mixing fraction (Figure 6A). However switching on the cation-π interactions at the weak cation-π limit diminishes such dynamic heterogeneity leading toward 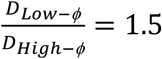 having weaker dependence on mixing fraction at Δϕ=2.08. At strong cation-π limit the dynamics are homogeneous throughout condensate. In absence of cation-π interactions and at intermediate heterogeneity of the sequence pairs (Δϕ=1.08), we have observed lower diffusivity of low-ϕ peptides to that of high-ϕ ones 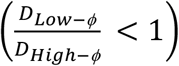 at *x*_*High−ϕ*=_ 0.25 indicating low-ϕ ones occupy the core and high-ϕ ones occupy the surface of the droplet (Figure 5B). Regardless of the cation-switching and mixing fraction, the chain reconfigurational timescale of high-ϕ peptides is 1.5 to 1.8 times longer than that of low-ϕ ones at intermediate heterogeneity of sequences (Figure 6B). The dynamic heterogeneity, 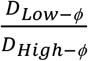 varies only up to 1.5 times at extreme *x*_*High−ϕ*_ in absence of sequence heterogeneity. (Figure 5C). Weak and strong cation-π interactions result in nearly homogenous condensates when the sequence heterogeneity of the pairs is relatively low or moderate (Δϕ=0.34-1.08). The invariance of, 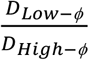 to variation of *x*_*High−ϕ*_ is introduced by both weak and strong cation-π limit. In absence of cation-π interactions, sequences with high-ϕ exhibit an average reconfiguration time that is 1.5 times faster than low-ϕ ones, particularly at low-to-intermediate *x*_*High−ϕ*_. Conversely, in presence of cation-π interactions, high-ϕ peptides exhibit similar dynamics to low-ϕ ones, which demonstrate tighter interaction through complementary hydrophobic and cation-π pairs and mixing (Figure 5B-C and 6C).

**Figure 5.**
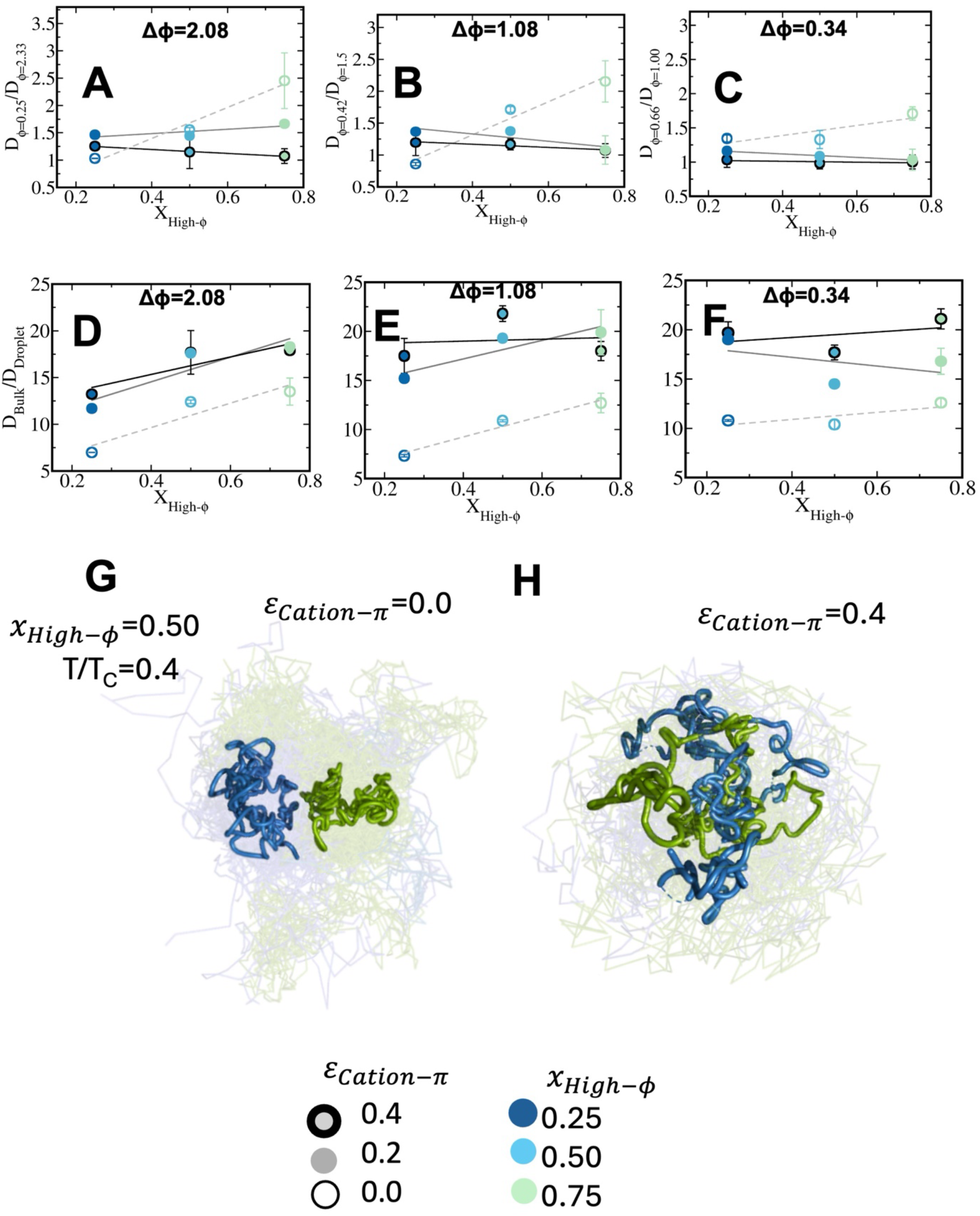
Translational dynamics in the condensate phase. Ratio of diffusivity among low and high-ϕ peptides in the dense phase 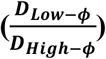 as a function of mixing fraction (*X*_*High*–ϕ_) at fixed T/T_C_=0.4 for designed systems having sequence heterogeneity (A) Δϕ = 2.08, (B) Δϕ = 1.08, and (C) Δϕ = 0.34. Peptide diffusivity in bulk relative to condensate phase 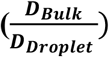 as a function of mixing fraction (*X*_*High*−ϕ_) at T/T_C_=0.4 for systems with (D) Δϕ = 2.08, (E) Δϕ = 1.08, and (F) Δϕ = 0.34 at various *X*_*High*−ϕ_. Colour codes represent mixing fractions (*X*_*High*−ϕ_). (G) Representative visual trajectories of peptides in condensate phase at extreme sequence heterogeneity (Δϕ = 0.5) in absence and presence of cation-π interactions at equimolar composition. Blue and green colours represent trajectories of peptides with low and high-ϕ values. In each panel, diffusivity ratios has been compared along cation-π strength ε_*Cation*−π_= 0. 0 (void symbol), ε_*Cation*−π_= 0. 2 *kcal*/*mol* (filled symbol) and ε_*Cation*−π_= 0. 4 *kcal*/*mol* (filled symbol with outer lines). While dashed lines showcase fits for systems without cation-π interactions, gray lines show fits for cation-π strength ε_*Cation*−π_= 0. 2 *kcal*/*mol* and black lines show fits for ε_*Cation*−π_= 0. 4 *kcal*/*mol*

**Figure 6.**
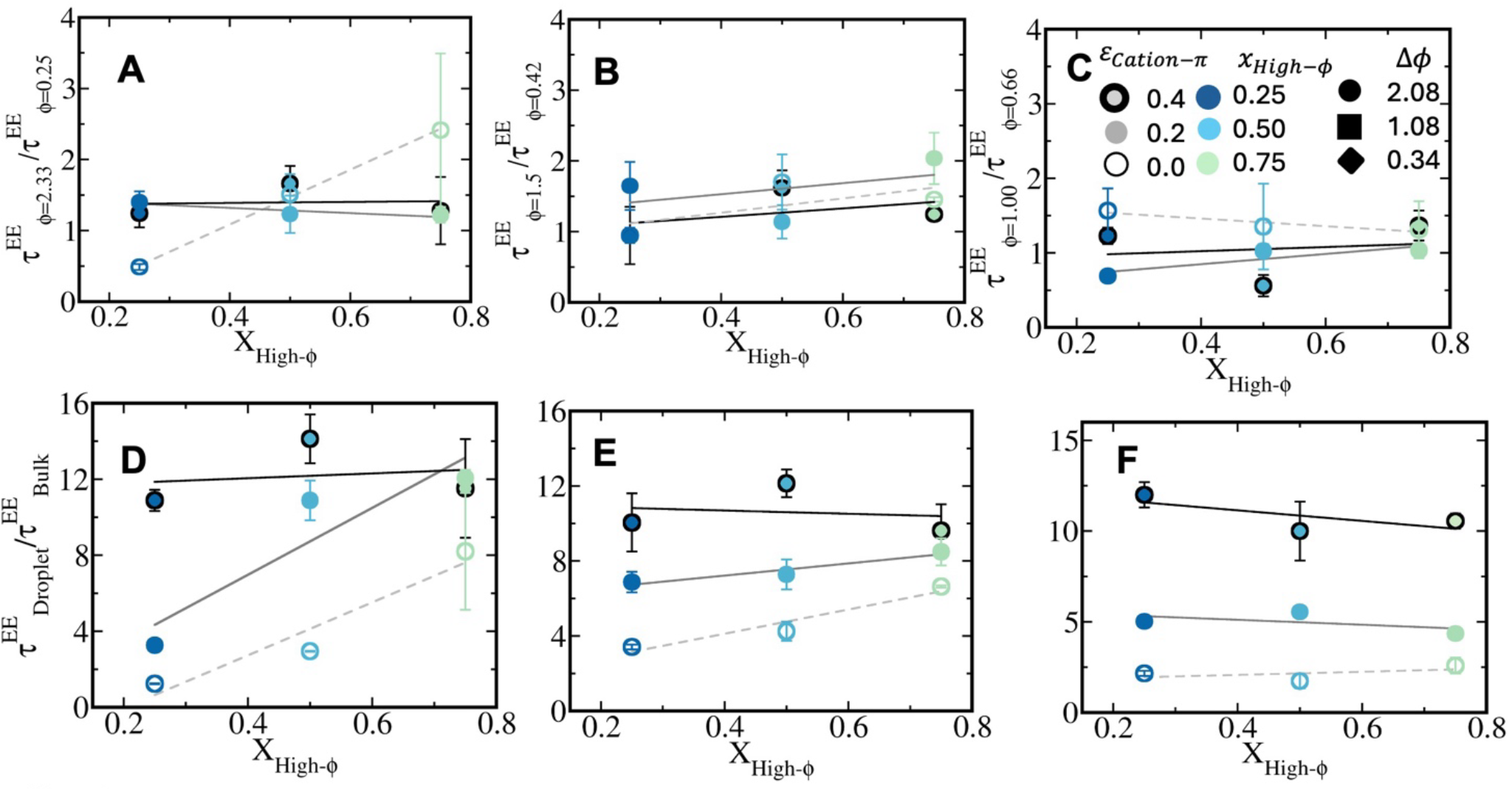
Chain reconfiguration dynamics in heterotypic condensates. Ratio of Reconfiguration lifetime 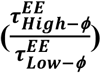 among low and high-ϕ peptides in condensate phase as a function of mixing fraction (*X*_*High*−*ϕ*_) for heterotypic systems with (A) Δϕ = 2.08, (B) Δϕ = 1.08, and (C) Δϕ = 0.34, respectively at T/T_C_=0.4. Average chain reconfiguration lifetime in the condensate phase compared to the bulk phase 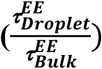 as a function of temperature for heterotypic systems with (D) Δϕ = 2.08, (E) Δϕ = 1.08, and (F) Δϕ = 0.34 respectively. Colour codes represent mixing fractions (*X*_*High*−*ϕ*_). In each panel, average reconfiguration timescale ratios have been compared along cation-π strength ε_*Cation*−π_= 0. 0 (void symbol), ε_*Cation*−π_= 0. 2 *kcal*/*mol* (filled symbol) and ε_*Cation*−π_= 0. 4 *kcal*/ *mol* (filled symbol with outer lines). While dashed lines showcase fits for systems without cation-π interactions, Gray lines show fits for cation-π strength ε_*Cation*−π_= 0. 2 *kcal*/ *mol* and black lines show fits for ε_*Cation*−π_= 0. 4 *kcal*/*mol*

Representative trajectories (cartoon) of low and high-ϕ peptide classes have been shown in all peptide trajectories (shown with transparent ribbons) in largest cluster showcasing significant de-mixed regions with differential diffusivity in absence of cation-π and a well-mixed condensate with homogeneous dynamics in presence of cation-π quantified in Figure 5(G-H).

To capture liquid-like nature of dense phase, we have computed average diffusivity of condensates and compared with the bulk diffusivity. For each system, a single polymer has been simulated in the exactly same conditions (temperatures) as that of the condensates in order to compute the diffusivity of the peptides in dilute phase. In absence of cation-π interactions, for extreme heterogeneity of the sequence pairs (Δϕ=2.08, Figure 5D), condensate diffusivity is nearly 7-times slower than dilute bulk at higher enrichment of sequences with charged residues (*x*_*High−ϕ*_= 0.25). Average reconfiguration timescale in dense phase varies in a correlated manner along translational diffusion in condensates. Reconfiguration timescale is nearly 1.5 times slower than bulk at extreme heterogeneity of designed peptides at low *x*_*High−ϕ*_. At extreme dominance of high-ϕ peptides, the slowdown enhances up to 6-times. At weaker cation-π limit 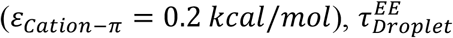 showcase an increment of 4-fold than 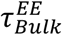 at low *x*_*High−ϕ*_ limit while at extremely high *x*_*High−ϕ*_, limit the slowdown marks as nearly 12-times with respect to bulk. At considerably high cation-π interaction strength, 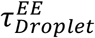 is invariant with respect to 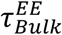 along mixing fraction and marks an uniform 12-times slowdown than bulk reconfiguration timescale of peptides. (Figure 6D)

Condensate diffusivity decreases with enhancement of sequences having higher hydrophobic content leading to 12-times slowdown at extreme mixing fraction of sequence with relatively higher hydrophobicity to charge ratio. Once we switch-on cation-π interactions, bulk diffusivity is nearly 15-18 times higher than condensates leading to 1.5 times rise than systems in absence of cation-π at all mixing fractions (Figure 5D). 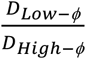 enhances as mixing fraction of peptides with enriched hydrophobicity increases. At low-to-intermediate sequence heterogeneity (Figure 5E-F), switching cation-π interactions at weak interaction limit causes 15-17 times slowdown of droplet diffusivity than bulk and such slowdown enhances with increase of *x*_*High−ϕ*_. At enhanced cation-π limit droplet diffusivity is nearly 20-times slower than bulk and invariant of mixing. In absence of cation-π interactions, at intermediate sequence heterogeneity 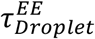 showcase 4-6 times enhancement of average reconfiguration timescale of peptides in dense phase as mixing fraction of peptides having high hydrophobic to charge ratio increases in the designed systems (Figure 6E). Switching on cation-π interactions at weaker limit enhances the slowdown of reconfiguration up to 8-fold and mildly dependent on the mixing fraction of the peptides. While cation-π interactions at stronger limit enhances slowdown of 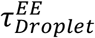 even more and suppresses any dependence of such timescale on peptide composition (Figure 6E). At lowest heterogeneity of pairs average reconfiguration timescale in droplet is 3-times slower than bulk in absence of cation-π interactions, however showcases 5-times slowdown at weaker cation-π limit and nearly 13 times slowdown at strong cation-π limit (Figure 6F) independent of mixing fraction.

### Condensate Architecture and Neighbour Exchange Dynamics

Condensate sphericity is highly dependent on cation-π switching in condensates. To decode condensate architecture, we have defined droplet anisotropy of largest cluster as follows:

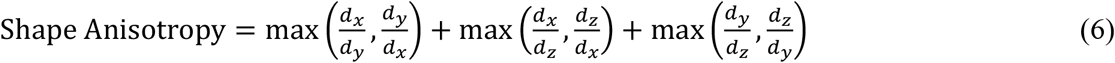

Such analysis showcases how much condensates closely resemble a sphere or deviates from the same. Ideally a condensate tends to be spherical to minimize the surface while inherent de-mixing or dominant electrostatic interactions and their exchange causes them to have deviation from spherical shape. In absence of cation-π interactions droplet shape anisotropy scaled to largest cluster is low enough at extremely low and high mixing fraction of sequences having high hydrophobicity to charged ratio (*x*_*High−ϕ*_), and enhances along peptide sequence heterogeneity (Figure 7 panels A-C). While we switch-on cation-π interactions, we have observed a decrease in shape anisotropy of condensates along sequence heterogeneity (Figure 7A-C,solid lines). We have observed the scaled anisotropy has enhanced at Δϕ=1.08 and 2.08 along cation-π strength significantly than other Δϕ variant at low *x*_*High−ϕ*_ limit due to strong electrostatic repulsion among like charged residues while initiating tight packing. At extreme enrichment of sequences with enhanced hydrophobicity fraction (Figure 7C), one observes a tighter packing complemented by hydrophobic interactions at stronger cation-π limit. In absence of cation-π, anisotropy at moderate *x*_*High−ϕ*_ is fairly large showcasing multicomponent dense phase coexisting and have nearly no dependence on sequence heterogeneity. In presence of cation-π interactions, enhanced sequence heterogeneity decreases anisotropy nearly 1.5-2 fold depending upon cation-π strength.

**Figure 7.**
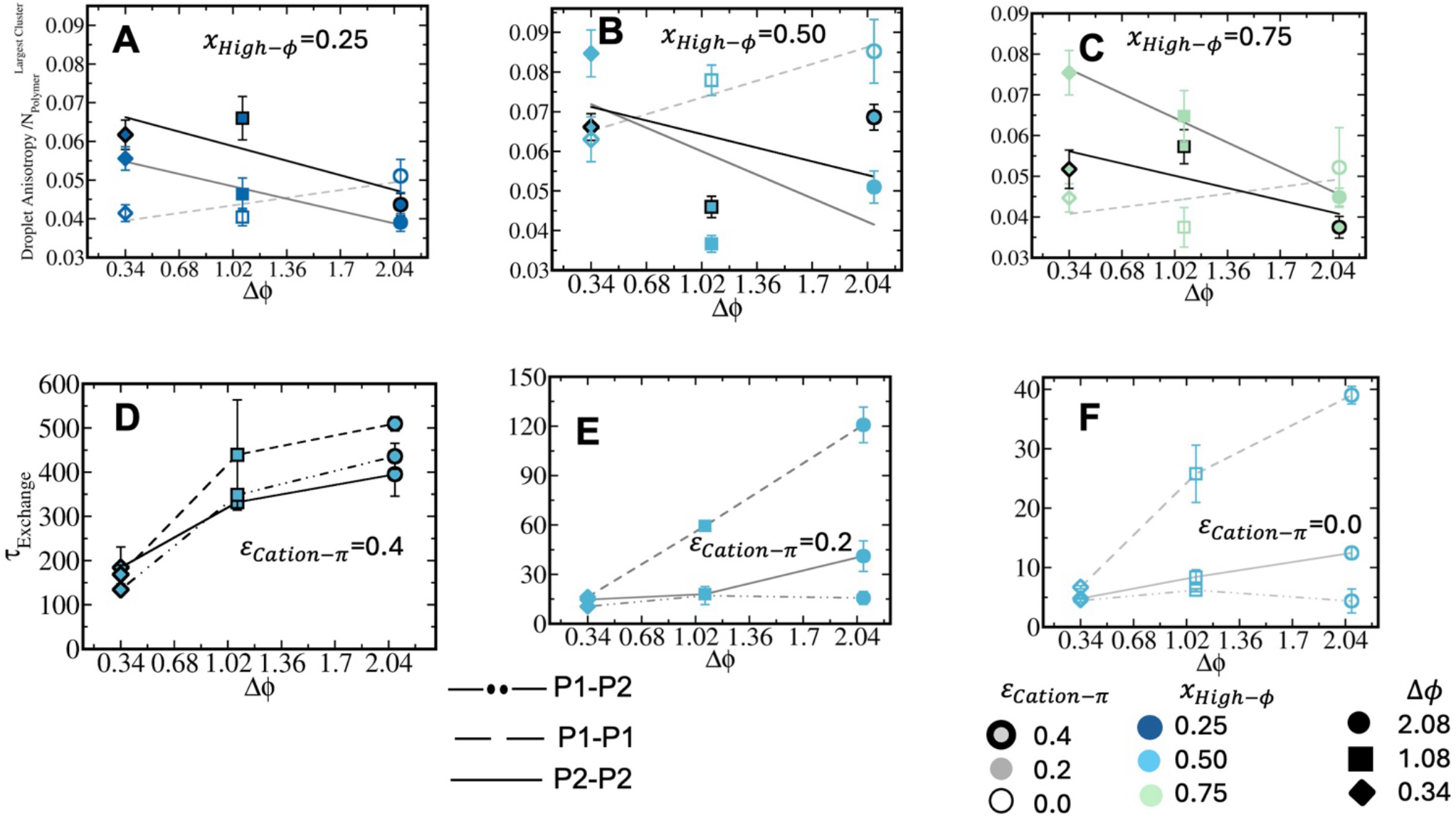
Droplet anisotropy scaled to cluster size as a function of sequence heterogeneity (Δϕ) for mixing fractions (A) *x*_*High*−ϕ_= 0. 25 (B) *x*_*High*−ϕ_= 0. 50, (C) *x*_*High*−ϕ_= 0. 75. Colour codes represent mixing fractions. Average neighbour exchange timescale (τ_Exchange_) for peptides as a function of the composition difference (Δϕ) for heterotypic systems at *X*_*High*−ϕ_= 0.50 along sequence heterogeneity of binary peptide pairs (Δϕ) at cation-π strength (D) 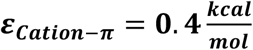, (*E*)ε_*Cation*−π_= 0. 2 *kcal*/*mol* and (E) ε_*Cation*−π_= 0. 0 *kcal*/*mol*. Circles, squares, diamonds and triangles represent Δϕ variants in descending order. Gray lines show fits for cation-π strength ε_*Cation*−π_= 0. 2 *kcal*/*mol* and black lines show fits for ε_*Cation*−π_= 0. 4 *kcal*/*mol*

In absence of cation-π interactions we have observed multiphase condensates with significant domain like architecture (Figure 2J), however once cation-π interactions are switched on, mixing of peptides is uniform in a condensates. One must note the effect of cation-π interactions are maximum when sequence heterogeneity is tremendously high. Similarly in absence of cation-π interactions, structural heterogeneity is also maximum at significant peptide heterogeneity. Such opposite effects cause inversion of a positive slope in droplet anisotropy along sequence heterogeneity to a negative one in presence of cation-π.

To elucidate multivalency induced exchange of sticker contacts in condensate phase which is intricately related to condensate sphericity, we have defined a time correlation function of sticker exchange as follows:

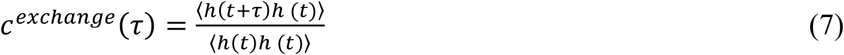

h(t) denotes a step function; h(t)=1 when two peptides are neighboring ones for a time frame and 0 otherwise. The correlation functions best fits to a bi-exponential function and the average neighbour exchange timescales for low-ϕ-low-ϕ (P1-P1), high-ϕ-high-ϕ (P2-P2) and cross contacts among low-ϕ-high-ϕ (P1-P2) has been plotted in Figure 7 (panels D-F) for equimolar abundance and at absolute temperature T=0.3 (below criticality) of the peptide pairs along sequence heterogeneity enhancement at different cation-π strengths. Until cation-π interactions are either absent or weak, P1-P1 and P2-P2 are relatively long lived and majorly at extreme sequence heterogeneity suggesting deficient mixing, while at strong cation-π limit a synergetic effect of P1-P2 cross sticker contacts mediated by cation–π interactions make both P1-P2 and P2-P2 interactions are fairly long lived. At weaker cation–π limit exchange of F-F and R-F are significant leaving the sticker exchange timescale for P2-P2 and P1-P2 low enough. One must note the comparison has been shown at an absolute temperature to turn-off the effect of temperature on the exchange kinetics (Figure 7(D-F),S8 and S9).

## Conclusion

In this work, we have investigated how cation–π interactions regulate stability, spatial organization, and internal dynamics of heterotypic peptide condensates in presence of sequence heterogeneity and compositional asymmetry. By systematically varying the hydrophobicity-to-charge mismatch between two peptide sequences, their relative mixing fraction, and the strength of cation–π interactions, we establish a molecular framework for understanding how an additional short-range cross-interaction can reshape the balance between self-association and heterotypic association within a condensed phase. Our simulations demonstrate that condensate organization cannot be understood from sequence heterogeneity or composition alone; rather, it emerges from the competition and cooperation among electrostatic, hydrophobic, and cation–π interactions.

In the absence of cation–π interactions, sequence heterogeneity produces pronounced spatial segregation within the condensate. The two peptide populations preferentially occupy different regions of the dense phase, giving rise to core–shell-like or domain-rich architectures whose orientation depends on the relative abundance of the two components. Peptides enriched in charged residues preferentially form the electrostatically stabilized core under charge-dominated conditions, whereas increasing the fraction of hydrophobic peptides shifts the organization toward a hydrophobic-rich core and a complementary outer region. Thus, composition does not simply alter the amount of each component within the condensate; it changes the dominant interaction network and consequently the spatial arrangement of the constituent molecules. This observation highlights the importance of considering composition and sequence heterogeneity as coupled variables when describing heterotypic phase separation.

The introduction of cation–π interactions substantially modifies this behaviour. Even at the weaker interaction strength, cation–π contacts increase condensate stability, while stronger cation–π interactions produce a considerably larger enhancement in the critical temperature and condensate density. The effect is particularly pronounced for highly heterogeneous peptide pairs, where increasing cation–π strength can substantially elevate the critical temperature and increase the density of the condensed phase. These results indicate that cation–π interactions provide an additional energetic pathway for condensation that can reinforce the cohesive interactions already present in the system. Importantly, this stabilization is not simply additive. The energetic analysis shows that cation–π interactions partially compensate for changes in electrostatic and hydrophobic contributions, revealing a redistribution of the interaction network as the composition and sequence mismatch are varied. At strong cation–π coupling, the energetic contribution from these interactions becomes comparable to, and in some conditions approaches, the magnitude of the principal electrostatic and hydrophobic contributions.

A particularly important consequence of introducing cation–π interactions is the conversion of spatially heterogeneous condensates into more uniformly mixed dense phases. In the highly heterogeneous systems, the absence of cation–π interactions leads to well-defined compositional domains, whereas increasing cation–π strength progressively erodes these spatial distinctions. This behaviour suggests that cation–π interactions effectively increase the compatibility between chemically distinct peptide populations by providing favourable cross-association pathways. In this sense, cation–π interactions function as a molecular bridge between otherwise competing interaction preferences: they allow charged and aromatic groups belonging to different peptide populations to associate directly, thereby reducing the thermodynamic advantage of segregation. The resulting condensate is consequently more compositionally homogeneous despite retaining substantial sequence heterogeneity.

The same coupling between interaction chemistry and organization is reflected in condensate dynamics. In the absence of cation–π interactions, translational diffusion and chain reconfiguration exhibit strong dependence on both sequence heterogeneity and composition. The peptide population occupying the more strongly interacting region of the condensate displays slower molecular motion, consistent with the formation of a dynamically distinct dense domain. As cation–π interactions are introduced, this dynamic contrast is progressively reduced, and the dependence of molecular mobility on mixing fraction becomes weaker. At sufficiently strong cation–π interactions, the reconfiguration dynamics become comparatively insensitive to composition, indicating that the additional cross-interaction homogenizes not only the spatial distribution of the peptides but also their local dynamical environment.

At the same time, the formation of stronger cation–π networks does not make the condensate dynamically faster. Rather, enhanced cohesive interactions substantially slow translational diffusion and internal chain reconfiguration relative to the dilute phase. The dense phase therefore becomes more dynamically constrained as the cation–π interaction strength increases.

This behavior is especially evident for highly heterogeneous systems, where strong cation–π interactions produce a pronounced slowdown while simultaneously suppressing the compositional dependence of the dynamics. Thus, cation–π interactions establish a notable relationship between molecular mixing and dynamical confinement: improved molecular compatibility and spatial homogenization are accompanied by stronger intermolecular coupling and slower molecular rearrangement.

The structural analysis further supports this picture. Without cation–π interactions, increasing sequence heterogeneity promotes deviations from spherical morphology and the emergence of domain-like internal organization. Upon introducing cation–π interactions, this trend is reversed, with increasing interaction strength generally reducing condensate anisotropy and producing more compact and homogeneous structures. At high cation–π strength, hydrophobic interactions can further cooperate with cation–π contacts to promote tighter packing, particularly in systems enriched in hydrophobic sequences. The resulting inversion in the dependence of condensate anisotropy on sequence heterogeneity demonstrates that a relatively modest change in the interaction network can qualitatively alter the relationship between molecular sequence and mesoscale morphology.

Taken together, these results establish cation–π interactions as an important regulatory interaction in heterotypic condensates. Their role extends beyond simply increasing the overall cohesive energy of the dense phase. Instead, cation–π interactions simultaneously influence phase stability, molecular partitioning, condensate morphology, translational diffusion, and conformational reconfiguration, thereby linking thermodynamic and dynamical properties of the condensate. The strongest effects arise when sequence heterogeneity is large, because this provides the greatest contrast between the self-association preferences of the two peptide populations and, consequently, the greatest opportunity for cation–π-mediated cross-association to reshape the interaction network. Conversely, when sequence heterogeneity is small, the condensate properties become less sensitive to composition because the two components already possess more similar interaction characteristics.

More broadly, our results suggest that the molecular organization of heterotypic condensates can be viewed as a tuneable balance between de-mixing interactions and cross-associative interactions. Sequence heterogeneity provides a driving force for spatial differentiation, whereas cation–π interactions introduce an interaction channel that favours molecular mixing. Composition determines how frequently these competing interactions can be realized and therefore controls the resulting balance between segregated and mixed states. The resulting condensate state is consequently not determined by a single interaction parameter but by the collective organization of multiple weak interactions operating over different spatial ranges. This perspective provides a useful framework for interpreting how sequence variation can be translated into distinct mesoscale condensate states.

The present study therefore demonstrates a general molecular principle: additional short-range cross-interactions can convert sequence heterogeneity from a source of spatial and dynamic segregation into a tuneable determinant of condensate stability and organization. By controlling the strength of cation–π interactions, it is possible to modulate whether chemically distinct peptide populations preferentially segregate into domains or form a dynamically coupled, homogeneous condensed phase. Such interaction-level control provides a potential route for understanding and ultimately engineering the internal organization and material properties of multicomponent biomolecular condensates. Future studies incorporating additional interaction motifs and more complex sequence architectures could extend this framework toward condensates containing multiple chemically distinct components and help establish predictive principles for how sequence chemistry, interaction specificity, and molecular composition collectively determine condensate structure and dynamics.

## Notes

### Competing Interest Statement

The authors have declared no competing interest.

### Summary of Updates

Figure caption of Figure 2 has been updated. A link to supporting figure S3 has been made in page 14

### Funding

- ANRF: ANRF/ECRG/2024/000734/LS
- IIT Jodhpur: I/SEED/MKZ/20230169

