## Supporting Information for "To mix or to de-mix? : Cation-π suppresses heterogeneity in local structure and dynamics of heterotypic Condensates"

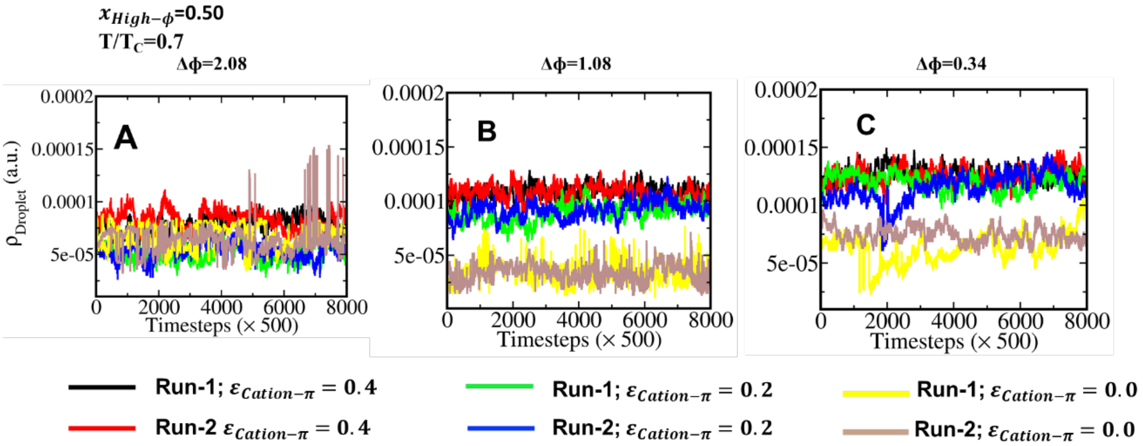


**Figure S1.** Convergence of condensate density obtained from two independent simulation runs for systems with different sequence-pair heterogeneities. The condensate density $\rho$, is shown as a function of simulation time for (A) $\Delta\phi$=2.08, (B) $\Delta\phi$ = 1.08, and (C) $\Delta\phi$ = 0.34, respectively. In each panel, the density profiles are shown for three different cation-π interaction strengths, $\varepsilon_{\text{cation-}\pi}$ = 0.4, 0.2, and 0.0. Black and red curves represent Run 1 and Run 2, respectively, for $\varepsilon_{\text{cation-}\pi}$ = 0.4; green and blue curves represent Run 1 and Run 2, respectively, for $\varepsilon_{\text{cation-}\pi}$ =0.2; and yellow and light-brown curves represent Run 1 and Run 2, respectively, for $\varepsilon_{\text{cation-}\pi}$ =0.0. The agreement between independent simulation runs demonstrates the convergence and reproducibility of condensate density across different sequence-pair heterogeneities and cation-π interaction strengths.

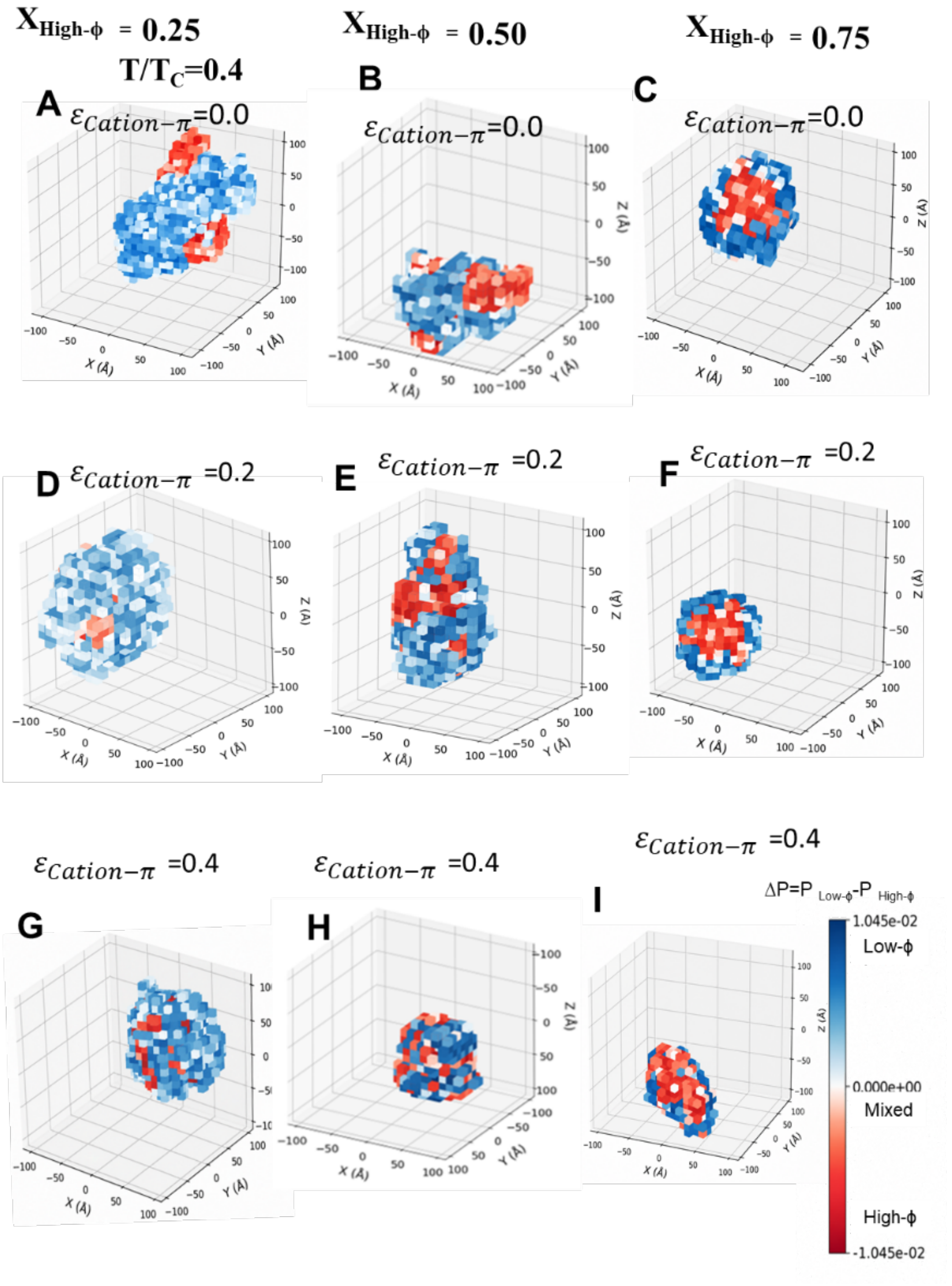


**Figure S2.** Representative three-dimensional probability order parameter $\Delta P = P_{Low-\phi} - P_{High-\phi}$ in largest cluster cubic grids illustrating the effect of mixing fraction on condensate heterogeneity for the Δϕ=2.08 system at different cation-π interaction strengths. Panels (A–C), (D–F), and (G–I) correspond to $\varepsilon_{\text{cation-}\pi}$ = 0.0, 0.2, and 0.4, respectively. Within each row, panels from left to right correspond to the mixing fractions $x_{High-f}$ = 0.25, 0.50, and 0.75, respectively. Red and blue regions represent high-$\phi$ and low-$\phi$ peptides, respectively. In the absence of cation-π interactions, the largest clusters exhibit pronounced spatial segregation of high-$\phi$ and low-$\phi$ peptides, whereas increasing cation-π interaction strength promotes more homogeneous mixing of the two peptide types within the condensate. The snapshots illustrate the dependence of condensate spatial organization and heterogeneity on mixing fraction and cation-π interaction strength.

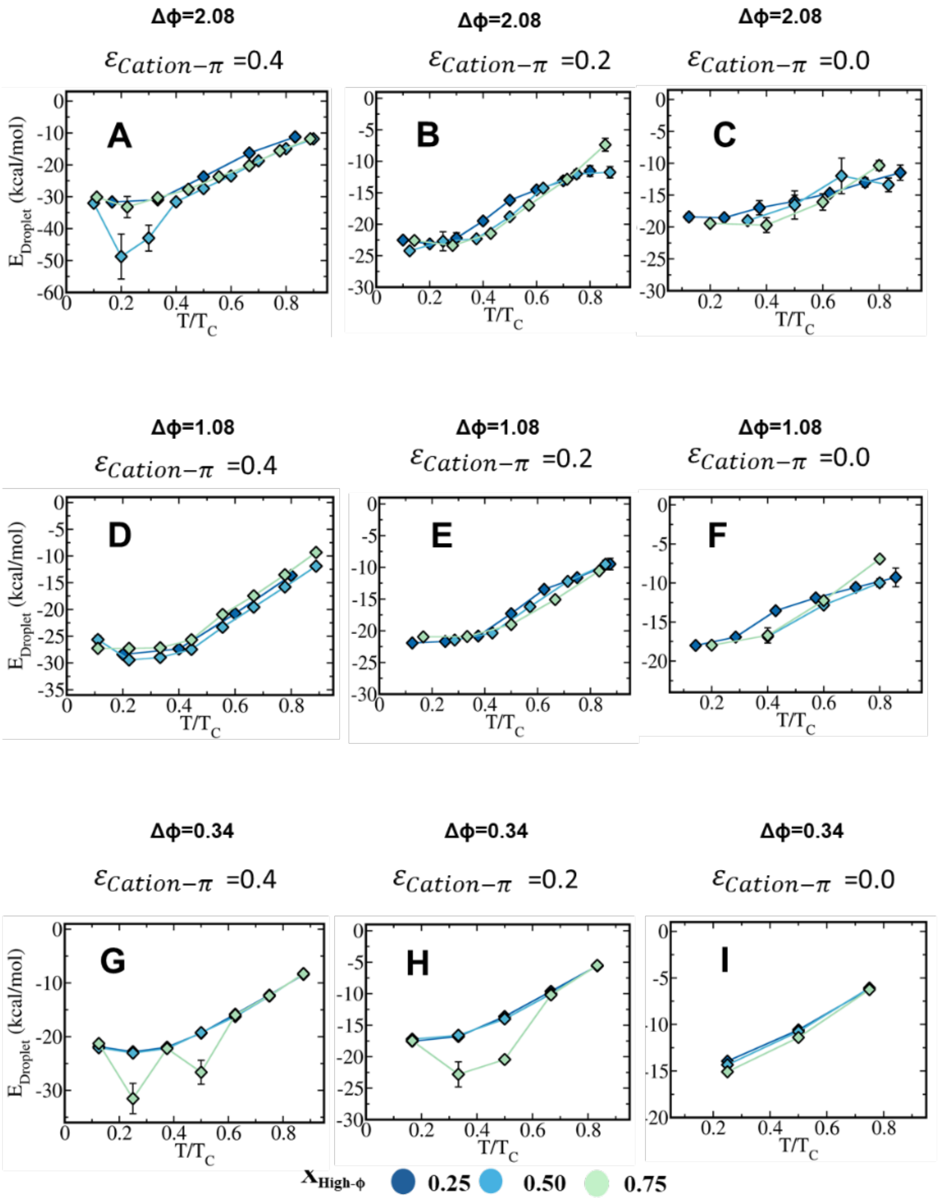


**Figure S3.** Temperature dependence of the average droplet energy, $E_{\text{droplet}}$, for different sequence-pair heterogeneities and cation-π interaction strengths. Panels (A–C) correspond to the Δϕ = 2.08 system, panels (D–F) correspond to the Δϕ = 1.08 system, and panels (G–I) correspond to the Δϕ = 0.34 system. Within each row, panels from left to right correspond to Δϕ $\varepsilon_{\text{cation-}\pi}$ = 0.4, Δϕ $\varepsilon_{\text{cation-}\pi}$ = 0.2, and without cation-π interactions, respectively. Thus, panels C, F, and I represent systems without cation-π interactions. The droplet energy is shown as a function of reduced temperature, T/T$_C$, for different mixing fractions, $x_{High-\phi}$ = 0.25, 0.50, and 0.75, represented by dark-blue, light-blue, and light-green symbols, respectively. The connecting lines guide the variation of droplet energy with temperature for each mixing fraction, while the error bars indicate the uncertainty in the calculated average droplet energies.

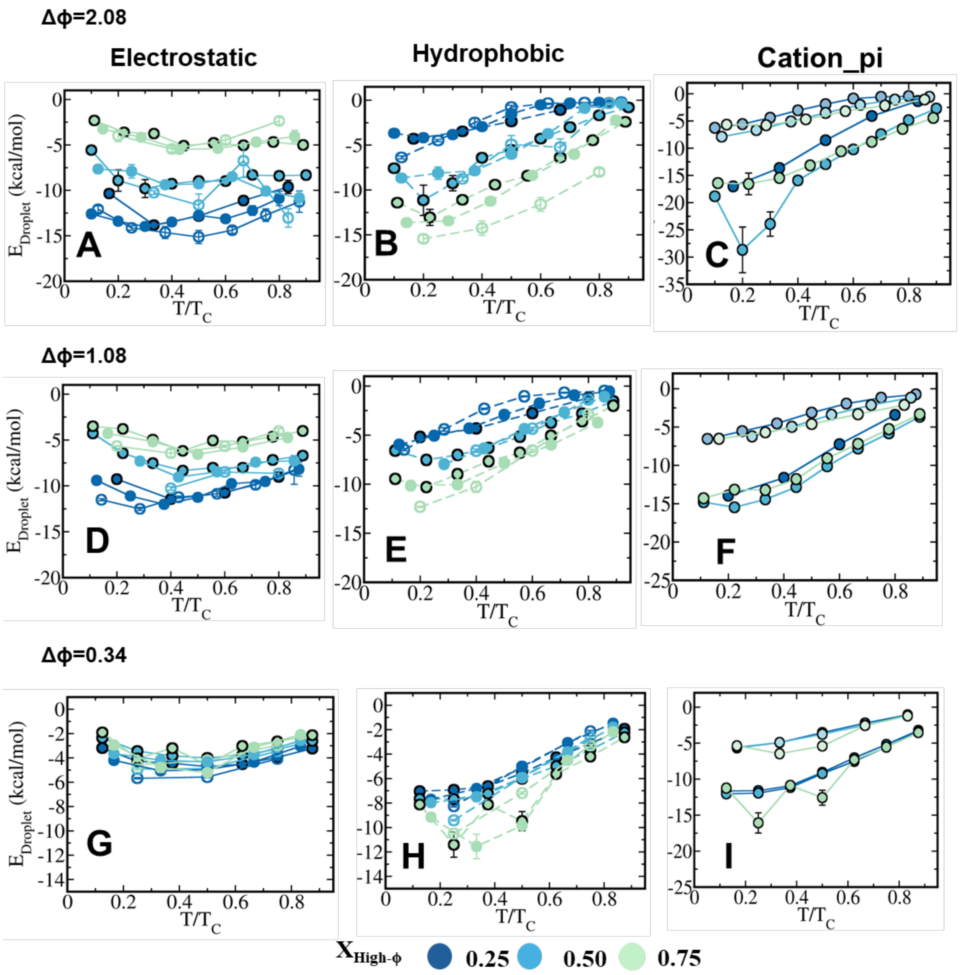


**Figure S4.** Temperature-dependent average droplet energy contributions from electrostatic, hydrophobic, and cation-π interactions for different sequence-pair heterogeneities at T/Tc = 0.4. Panels (A–C) correspond to the Δϕ = 2.08 system, panels (D–F) correspond to the Δϕ =1.08 system, and panels (G–I) correspond to the Δϕ = 0.34 system, respectively. Panels (A), (D), and (G) show the electrostatic energy, while panels (B), (E), and (H) show the hydrophobic energy, with results presented for $\varepsilon_{\text{cation-}\pi} = 0.4$, 0.2, and without cation-π interactions. Panels (C), (F), and (I) show the corresponding cation-π interaction energy for systems with cation-π interactions. The different symbols/colours correspond to mixing fractions $x_{High-\phi} = 0.25$, 0.50, and 0.75. Error bars represent the uncertainty in the calculated average energies. The figure illustrates the contributions of different interaction components to the droplet energetics at T/Tc=0.4 across different sequence-pair heterogeneities and mixing fractions.

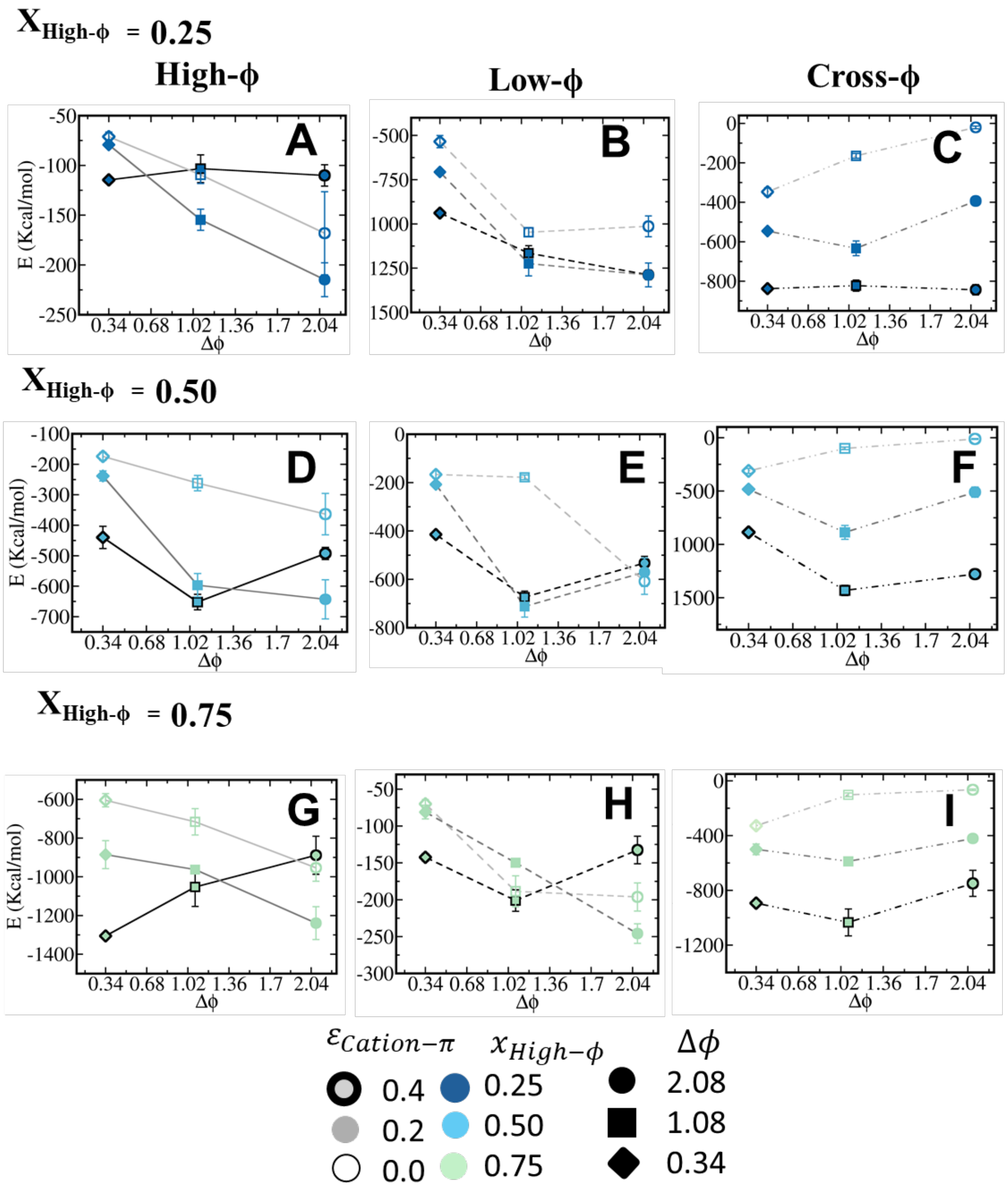


**Figure S5.** Dependence of the average interaction energy on sequence-pair heterogeneity, Δϕ at T/Tc=0.4 for different mixing fractions and cation-π interaction strengths. Panels (A–C), (D–F), and (G–I) correspond to $x_{High-\phi}$ = 0.25, 0.50, and 0.75, respectively. Within each row, panels from left to right show the interaction energies between high-$\phi$—high$\phi$ (P2-P2 ) low-$\phi$—low-$\phi$(P1-P1), and high-$\phi$-low-$\phi$ (P1-P2) peptide pairs at T/TC=0.4, respectively. Different symbol fillings represent cation-π interaction strengths $\varepsilon_{\text{cation-}\pi}$ = 0.4, 0.2, and 0.0, while different colours correspond to the mixing fractions $x_{High-\phi}$ = 0.25, 0.50, and 0.75. Different symbols represent the sequence-pair heterogeneities Δϕ=2.08, 1.08, and 0.34. Error bars indicate the uncertainty in the calculated average interaction energies. The figure illustrates the dependence of high-$\phi$, low-$\phi$, and cross-$\phi$ interaction energies on sequence-pair heterogeneity, mixing fraction, and cation-π interaction strength.

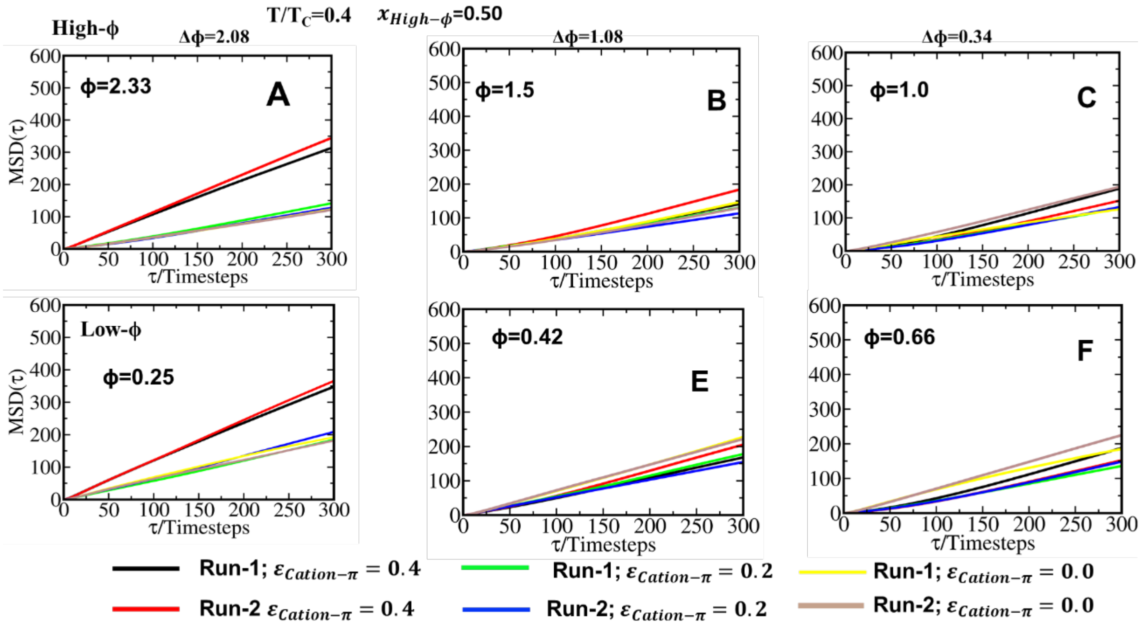


**Figure S6.** Representative mean-square displacement (MSD) profiles obtained from two independent simulation runs at mixing composition $X_{\text{High}-\phi} = 0.50$ and T/Tc=0.4. MSD profiles are shown for the designed systems with different sequence-pair heterogeneities, Δϕ = 2.08, 1.08, and 0.34. Panels (A-C) correspond to high-ϕ peptides, while panels (D–F) show the corresponding MSD profiles for low-ϕ peptides, respectively. For each system, MSD profiles are shown for three cation-π interaction strengths $\varepsilon_{Cation-\pi} = 0.2$, 0.2, and 0.0. Black and red curves represent Run 1 and Run 2, respectively, for $\varepsilon_{Cation-\pi}$= 0.4 green and blue curves represent Run 1 and Run 2, respectively, for $\varepsilon_{Cation-\pi}$= 0.2; and yellow and light-brown curves represent Run 1 and Run 2, respectively, for $\varepsilon_{Cation-\pi}$=0.0. The agreement between the two independent runs demonstrates the reproducibility of the chain dynamics across different cation-π interaction strengths and sequence-pair heterogeneities.

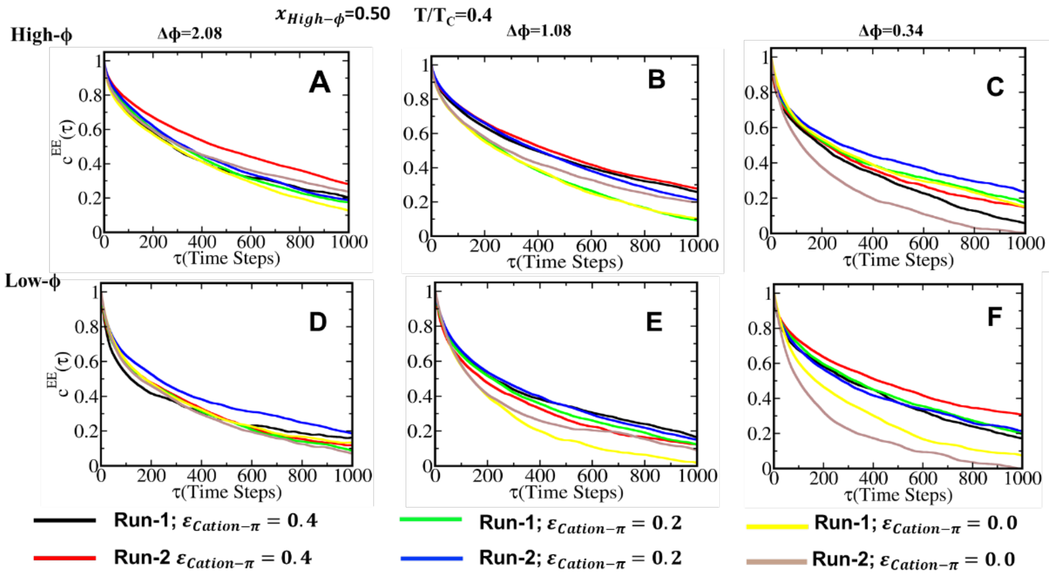

**Figure S7.** Representative end-to-end distance time correlation functions of high-ϕ and low-ϕ peptides in heterotypic condensates at T/Tc=0.4 and $x_{High-\phi} = 0.50$. Panels (A-C) correspond to the three sequence-pair heterogeneity systems with Δϕ=2.08, 1.08, and 0.34, respectively, while panels (D–F) show the corresponding results for low-ϕ peptides. The curves represent end-to-end distance time correlation functions obtained at different cation-π interaction strengths $\varepsilon$ =0.4, 0.2, and 0.0 kcal/mol from two independent simulation runs. Black and red curves represent Runs 1 and 2, respectively, for $\varepsilon_{Cation-\pi}$ =0.4 kcal/mol; green and blue curves represent Runs 1 and 2, respectively, for $\varepsilon$ =0.2; and yellow and light-brown curves represent Runs 1 and 2, respectively, for $\varepsilon$ =0.0. The results illustrate the dependence of chain reconfiguration dynamics on sequence-pair heterogeneity, peptide type, and cation-π interaction strength.

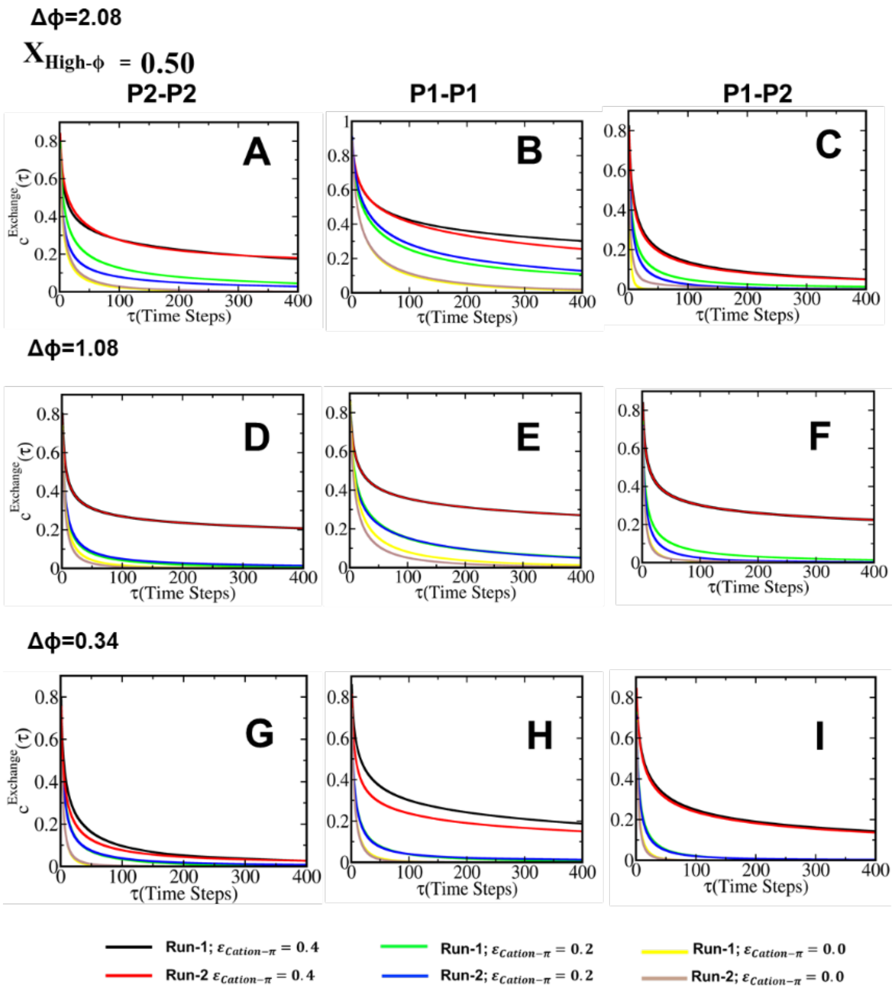


**Figure S8.** Representative neighbour contact exchange time correlation functions in the condensate phase at T/Tc=0.4 and $x_{High-f} = 0.50$. Panels (A–C), (D–F), and (G–I) correspond to the Δϕ = 2.08, Δϕ = 1.08, and Δϕ = 0.34 systems, respectively. Within each row, the panels correspond to high-$\phi$-high-$\phi$ (P2–P2), low-$\phi$ –low-$\phi$ (P1–P1), and low-$\phi$ –high-$\phi$ (P1–P2)

neighbour contacts, respectively. In each panel, black and red curves represent Run 1 and Run 2, respectively, for cation-π interaction strength $\varepsilon_{Cation-\pi}$ = 0.4; green and blue curves represent Run 1 and Run 2, respectively, for $\varepsilon_{Cation-\pi}$ = 0.2; and yellow and light-brown curves represent Run 1 and Run 2, respectively, for $\varepsilon_{Cation-\pi}$ = 0.0 The comparison of the exchange time correlation functions across different cation-π interaction strengths and sequence-pair heterogeneities illustrates the effect of cation-π interactions on the dynamics of different neighbour contact types within the condensates.

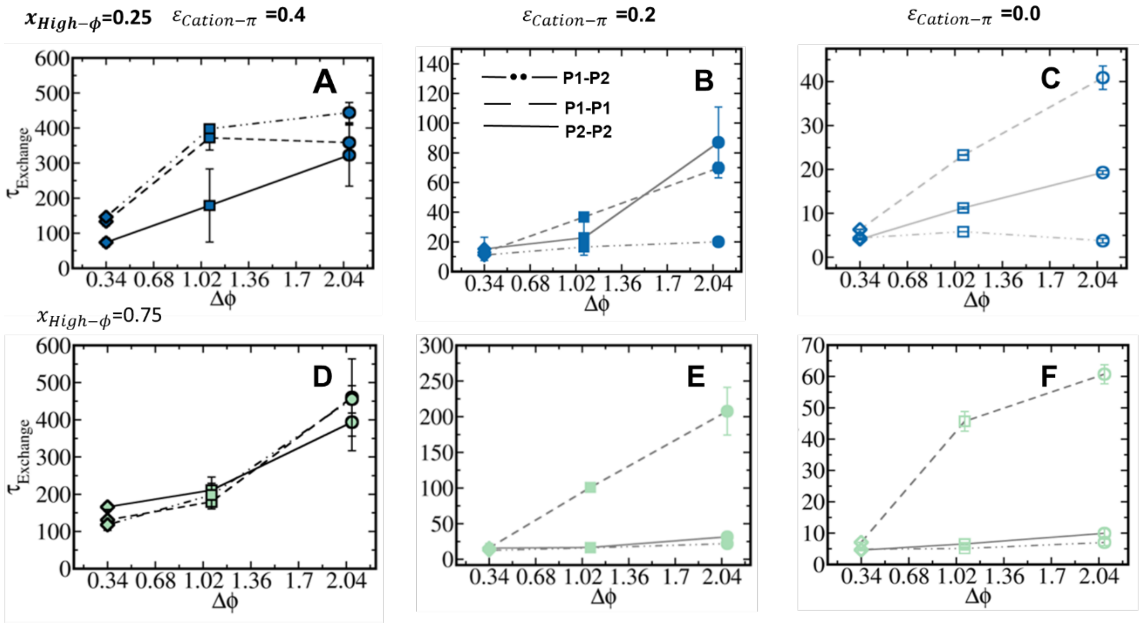


**Figure S9.** Dependence of the neighbour contact exchange time, $\tau_{\text{Exchange}}$, on sequence-pair heterogeneity Δϕ at different mixing compositions and cation-π interaction strengths. Panels (A–C) correspond to $x_{High-\phi}$= 0.25, while panels (D–F) correspond to $x_{High-\phi}$ = 0.75. Within each row, panels from left to right correspond to cation-π interaction strengths $\varepsilon_{\text{cation-}\pi}$ = 0.4, 0.2, and 0.0, respectively. Different symbols and line styles represent the exchange times of high-$\phi$–high-$\phi$ (P2–P2), low-$\phi$–low-$\phi$ (P1–P1), and cross (P1–P2) contacts, respectively. The corresponding error bars represent the uncertainty in the calculated exchange times. The figure demonstrates the dependence of neighbour contact exchange dynamics on sequence-pair heterogeneity, mixing composition, and cation-π interaction strength.
